# CapP mediates the structural formation of biofilm-specific pili in the opportunistic human pathogen *Bacillus cereus*

**DOI:** 10.1101/2024.02.27.582368

**Authors:** Ana Álvarez-Mena, Muhammed Bilal Abdul Shukkoor, Joaquín Caro-Astorga, Melanie Berbon, María Luisa Antequera-Gómez, Axelle Grélard, Brice Kauffmann, Estelle Morvan, Antonio de Vicente, Birgit Habenstein, Antoine Loquet, Diego Romero

**Author notes:** Lead contact X: @diegoromerohi (Diego Romero).

## Abstract

Polymeric proteinaceous filaments are structural scaffolds that diversify the functionality of the bacterial extracellular matrix. Here, we report a previously uncharacterized bacterial factor called *bc1280* that is exclusive to *B. cereus* group and indispensable for the establishment of a biofilm lifestyle. We propose that BC1280 is an essential chaperone for the assembly of the filamentous platform that tightly controls the polymerization of heteropili containing CalY and TasA as major subunits in a concentration-dependent manner. Additionally, BC1280 modulates the expression of EPS via an uncharacterized pathway that is activated by a protease and an ECF-type sigma factor. The pilus biogenesis system described in this work highlights the complexity of extracellular matrix assembly in *B. cereus* and introduces a singular three-part structuration mechanism during biofilm formation and maturation.

**Graphical abstract:** During the process of biofilm formation in *B. cereus*, TasA, CapP, and CalY serve as the main components, in addition to other factors, ensuring that the extracellular matrix correctly assembles. The three proteins are produced and processed by the signal peptidase SipW and subsequently secreted through the Sec pathway in an unfolded conformation. At early biofilm formation stages, CapP is initially present at low levels, associates with the cell wall and is secreted into the ECM where it interacts with unfolded subunits of CalY, facilitating CalY folding and initiating fibril growth. CalY serves as a nucleator for incorporating TasA subunits into the pilus, forming TasA-CalY heteropolymers. Once the ECM scaffold is established, CapP levels increase, forming stable complexes with CalY that prevent its folding and arrest filament growth.

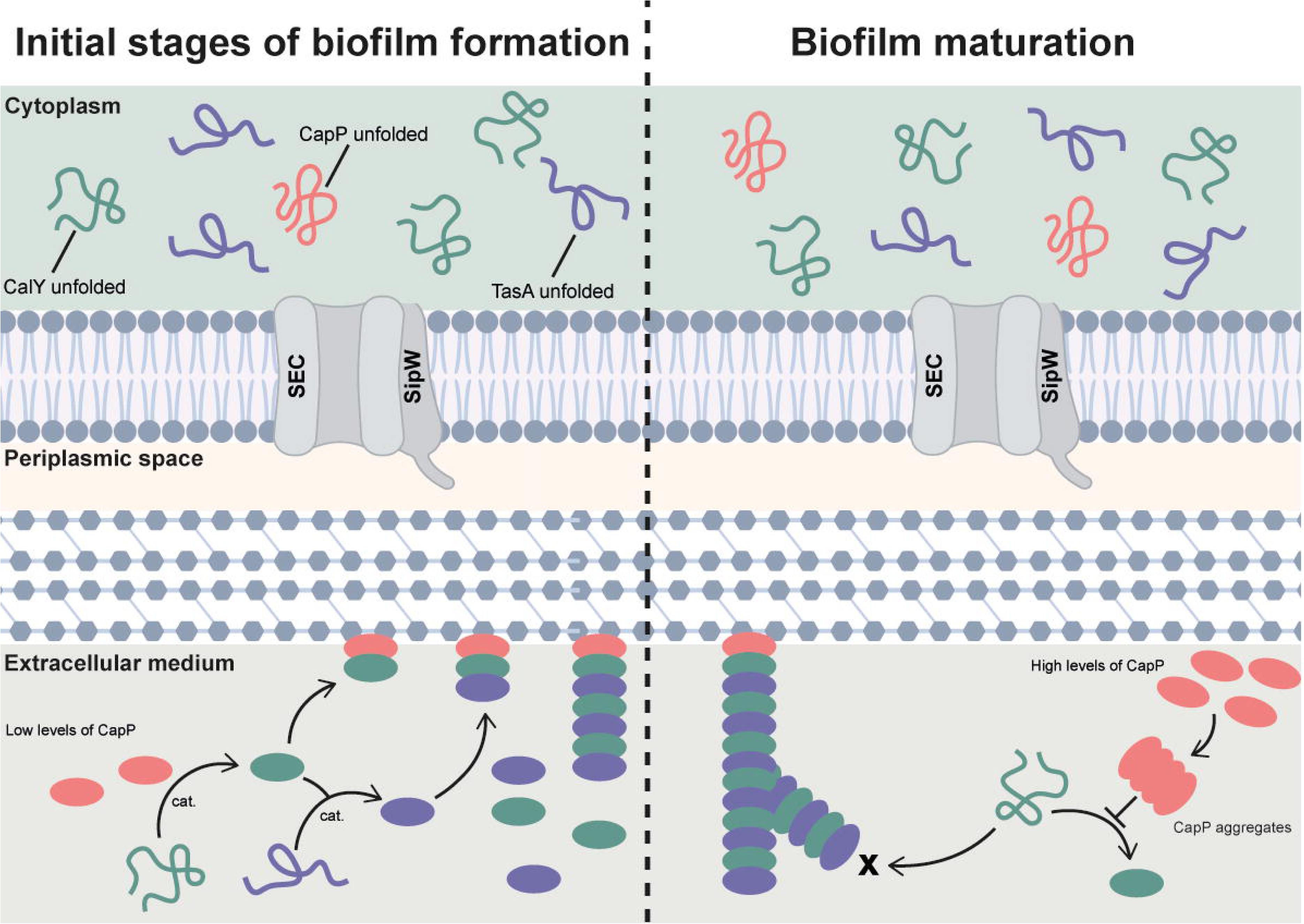

## INTRODUCTION

*Bacillus cereus*, a bacterium commonly found in soil, is capable of colonizing the gastrointestinal tracts of mammals or arthropods; *B. cereus* is contracted through the ingestion of contaminated food, including fresh vegetables and ready-to-eat meals.^1–4^ Consequently, *B. cereus* strains are frequently identified as the etiological agent of gastrointestinal diseases outbreaks, and the diseases can progress through emetic or diarrheal symptoms.^5,6^ As approximately 70% of microbial infections in humans are reportedly associated with biofilms, much effort has been dedicated to understanding the genetic basis and structural components that, upon activation, lead to the formation of these bacterial communities.^7,8^ The transition from a planktonic to a sessile lifestyle is metabolically focused on the synthesis of proteins, polysaccharides and extracellular DNA (eDNA), which ultimately organize the extracellular matrix (ECM).^9,10^ The ECM performs a wide range of functions, including adhesion to biotic and abiotic surfaces, regulation of nutrient and signaling molecule transport, defense against physical-chemical stress, and virulence, and provides support for biofilm architecture.^11,12^ One of the most attractive components of the ECM is the fibril-forming proteins that serve as structural scaffolds, facilitating the assembly of the microbial community and contributing to surface adhesion, among other biological roles.^13,14^ These proteins, termed functional amyloid-like proteins, share remarkable similarities with their pathogenic counterparts in humans, highlighted by their propensity to polymerize into highly ordered fibrillary aggregates, resistance to diverse environmental challenges, and their enrichment in β-sheet content.^15–18^

The development of *B. cereus* biofilms involves genetic pathways and structural components of the ECM identical to those found in *B. subtilis*, a closely related species. However, differences arise and are predicted to have ecological and pathological implications.^19,20^ Complementary reports have described the role of flagella, matrix proteins, eDNA and exopolysaccharides in the formation of biofilms on *B. cereus.*^21–24^ A previous study identified a specific genomic region in *B. cereus* that is orthologous to the corresponding region in *B. subtilis*, and it is expressed mainly in biofilm-associated cells.^25,26^ This region encodes an ortholog of *sipW;* two orthologs of *tasA*, called *tasA* and *calY*; the main biofilm regulators *sinI* and *sinR*; and an uncharacterized locus called *bc1280*. Interestingly, no orthologs for *tapA* are present in *B. cereus*. SipW is the signal peptidase that processes TasA and TapA for efficient secretion, and both TasA and CalY of *B. cereus* conserve the signal peptide with the canonical cleavage site recognized by the cognate peptidase SipW.^27,28^ The genes *sipW* and *tasA* constitute an operon, and *calY* is expressed independently.^25^ The deletion of *tasA* or *calY* resulted in differential defects in biofilm formation, suggesting that the product of each gene might be involved in cell-to-abiotic surface or cell-to-cell interactions, respectively.^25^ TasA from *B. subtilis* has been described as a globular protein in its monomeric state that polymerizes into filamentous structures, which can be amyloid or non-amyloid, depending on the experimental context.^29–33^ TasA native fibers, observed in vivo in biofilm samples and also in vitro upon extraction, exhibit an enrichment in β-sheet conformation while preserving the β-rich globular structure of the monomer. Consequently, these TasA monomers polymerize into filaments via a donor strand complementation mechanism, and as a result, their structure diverges from the canonical cross-β organization associated with amyloid proteins.^33^ Nevertheless, recombinant TasA purified by heterologous expression in *Escherichia coli* has been shown to form fibrils with typical amyloid properties, such as the cross-beta signature.^29,32^ TasA might therefore adapt its secondary and tertiary structure in a context-dependent manner, forming either an amyloid conformation or a β-rich globular fold. Extensive structural studies on TasA of *B. subtilis* exist, but only El-Mammeri et al. have reported singularities in recombinantly produced TasA and CalY from *B. cereus,* based on their secondary structure propensity, despite the cross-β amyloid pattern displayed by these paralogous proteins.^32^

In this study, we provide the first evidence that the *bc1280* product is indispensable for the growth of *B. cereus* biofilms, relying on two complementary functions. The first is a structural role, i.e., the promotion of CalY polymerization, which is related to the correct polymerization of TasA-CalY heterofibrils, essential for the formation and stability of the ECM scaffold. The second is a regulatory function in which the expression of biofilm-related EPSs is controlled via an undescribed sigma-antisigma pathway. Moreover, our findings indicate that when BC1280 is expressed heterologously in *E. coli*, the N-domain displays a predominantly β-sheet conformation within its rigid core, while the tandem repetitions found in the C-domain potentially facilitate the interaction with CalY. From its localization in the cell wall, BC1280 defines and triggers the polymerization of CalY, which in turn incorporates TasA into nascent heterofilaments. Based on these findings, BC1280 was renamed CapP, which is a CalY-assisting polymerization protein. Our study provides a new paradigm for the complex assembly of this proteinaceous appendage in *B. cereus* biofilms, identifies CapP as a structural orchestrator for efficient ECM development, and highlights CapP as a promising target for precisely targeted antibiofilm therapies.

## RESULTS

### *bc1280* is an essential locus in the developmental program of biofilm formation

First, a phylogenetic analysis of BC1280 was performed using the amino acid sequence of the type strain *B. cereus* ATCC14579 as a reference. The results revealed that BC1280 was present within the *B. cereus* group but not within the phylogenetically related species *B. subtilis* (Figure S1A). The *bc1280* product was annotated as a hypothetical protein with the following putative regions (Figure 1A): i) a signal peptide from position 1 to 39 predicted to be processed by a signal peptidase type I (SignalP5.0) (Figure S1B); ii) an N-terminal domain from position 40 to 189 (the N-domain from now on) that contains a highly conserved domain of unknown function (from I^33^ to E^157^) annotated as DUF4047; and iii) a C-terminal domain (the C-domain from now on) from position 190 to the C-terminal end of the protein. This domain is characterized by a variable number of perfect repetitions of QKKV/AEE motifs, depending on the *B. cereus* strains (Figure S1C).

**Figure 1.**
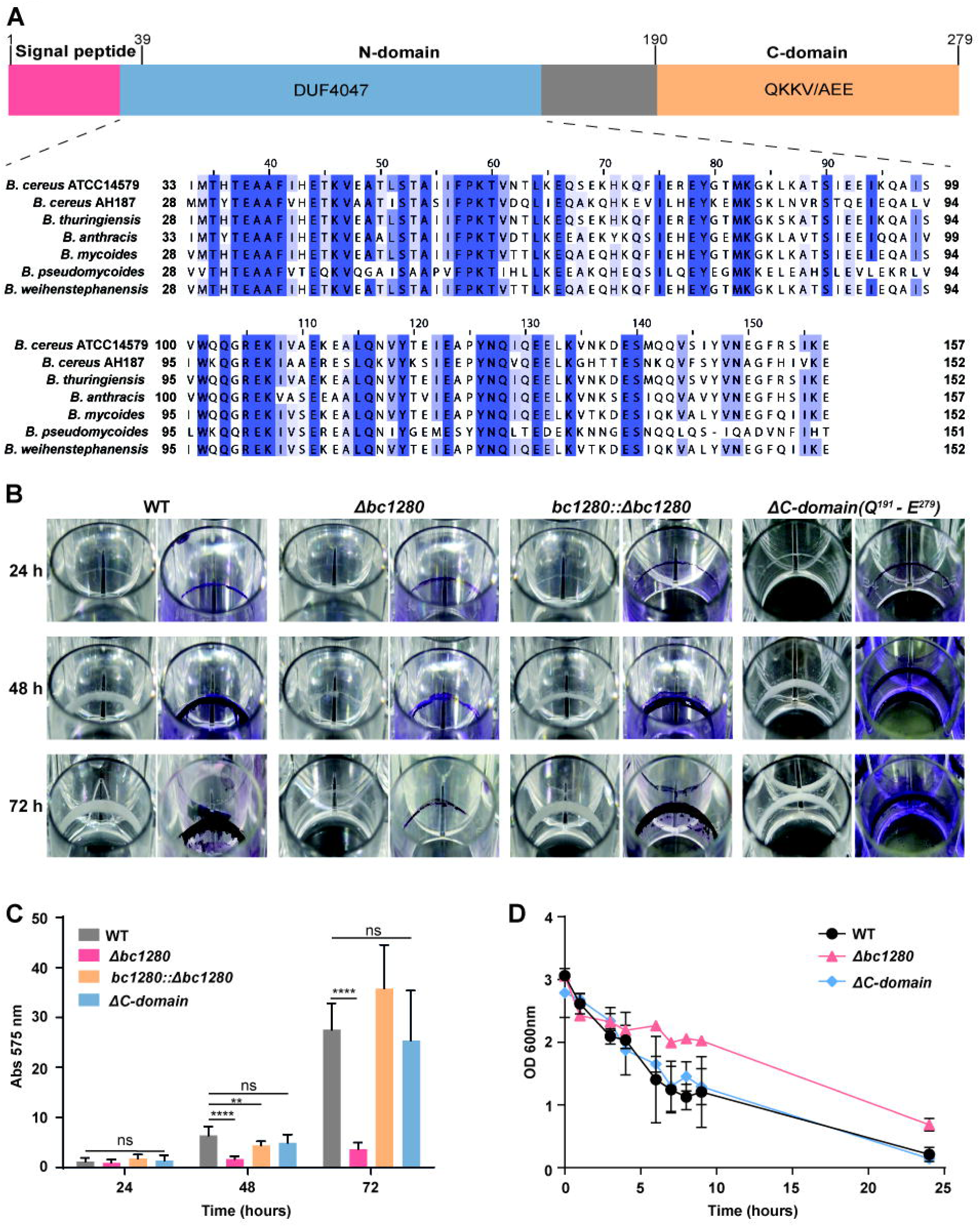
*bc1280* is an essential locus in the developmental program of biofilm formation. **A)** Schematic domain composition of BC1280. The signal peptide (M1-A39) is shown in red; the domain of unknown function, DUF4047 (I33-E157), which is located within the N-domain, is highlighted in blue; and the C-domain containing the repetitive region (Q190-E 279) is shown in yellow. The amino acid alignment of DUF4047 among different strains of the *B. cereus* group revealed that it is highly conserved, with the residues that are 100% conserved in all the strains marked in color. The color pattern is determined by the amino acid conservation. **B)** Biomasses formed by the *B. cereus* ATCC14579 wild-type strain and the mutants that adhered to abiotic surfaces were evaluated at different time points. Biofilms were stained with crystal violet to enable visualization and subsequent quantitative comparison. **C)** The amount of biofilm mass for each strain was estimated. Three independent experiments were conducted, each with three replicates. Average values are shown, and the error bars represent the standard deviation for each strain. Multiple comparisons were performed at each time point, with the wild-type serving as the control, and ordinary one-way ANOVA was used (** p value < 0.01 and **** p value < 0.0001). **D)** Auto-aggregation kinetics due to cell-to-cell interactions were studied for the wild-type, *Δbc1280* and *ΔC-domain* strains. The optical density at 600 nm at the air-liquid interface of the cell suspensions was measured each hour. The initial optical density was set to 3 at the start of the experiment, and the samples were incubated without agitation at room temperature. The error bars indicate the SDs calculated from three biological replicates.

A previous transcriptome analysis of the genetic features that distinguish planktonic and biofilm cells in *B. cereus* showed that *bc1280* is specifically expressed in biofilm-associated cells, and a complementary study revealed that the expression of *bc1280* comprised the regulon of the biofilm-specific master regulator SinR.^34^ We confirmed the biofilm-specific expression of *bc1280* via RT‒qPCR analysis of samples taken at 48 hours (Figure S1D) and determined that *bc1280* is transcribed as a monocistronic unit independent of the previously reported transcriptional units *sipW-tasA* or *calY* (Figure S1E).^25^ The biofilm of the *B. cereus* wild-type strain ATCC14579 was characterized by a white biomass ring that adhered to the plate wells; this ring was usually visible at 24 hours and reached maturity at 72 hours (Figure 1B). A *bc1280* knockout mutant strain (*Δbc1280*) was completely arrested during biofilm formation, a phenotype that differed from the less adhesive or thinner biomass of *ΔtasA* or *ΔcalY,* respectively (Figure 1B).^25^ According to the distribution and conservation observed in our phylogenetic analysis, deletion of *bc1280* in the emetic strain of *B. cereus* AH187, which belongs to a different *B. cereus* subgroup, resulted in the complete elimination of pellicle formation at the air‒liquid interface compared to that in the wild-type strain (Figure S1F).

The wild-type phenotype for biofilm formation was completely restored through complementation of *Δbc1280*, which was achieved through ectopic integration of the *bc1280 locus* (*bc1280*::Δ*bc1280*) under the control of its own putative promoter at a neutral locus (Figure 1B). Moreover, these findings suggested that the promoter for the expression of *bc1280* must be contained within the noncoding sequence between the open reading frame of *tasA* and *bc1280*. BC1280 contains a distinctive C-domain with perfect tandem repetition. As previously described, these characteristic regions often have significant structural and functional importance for the stabilization of protein folding overall.^35^ Therefore, we investigated whether the C-domain is involved in biofilm formation. The biofilm formation phenotype of *Δbc1280* was still rescued by a version of BC1280 that lacked the C-domain from Q^191^-E^279^ (Δ*C-domain*) (Figure 1B). Quantitative measurements at 575 nm of the biomass stained with crystal violet reinforced the visual differences in biofilm thickness (Figure 1C). No significant differences were observed between the strains at 24 hours, when initial cell attachment precedes biofilm maturation; however, differences became evident at 48 hours, coinciding with the initiation of ECM assembly and the growth of biofilms. After cells are initially attached to surfaces, cell-to-cell interactions occur that facilitate the subsequent growth of the biofilm, a cellular phenomenon that can be monitored through auto-aggregation experiments. The wild-type and *ΔC-domain* strains exhibited sharp aggregation kinetics; in contrast, *Δbc1280* did not form sediments at the bottom of the tube at the end of the 24-hour experiment (Figure 1D). Taken together, these results indicate that the product encoded by *bc1280* is involved in the developmental stages of bacterial cell-to-cell adhesion and the subsequent assembly and maturation of the ECM, a function that may rely on the N-domain of the protein.

### The biofilm-defective phenotype of *bc1280* is independent of the expression level of *tasA* or *calY*

The *Δbc1280* phenotype may result from changes in the expression levels of ECM components or structural malfunctioning in the polymerization of TasA-CalY heterofilaments; these two alternative explanations are not mutually exclusive. A whole-transcriptome analysis of the *Δbc1280* strain revealed 35 genes that were overexpressed and 4 genes that were repressed compared to those in the wild-type strain at 24 h. However, none of these genes were predicted to be associated with biofilm formation (Table S1). Notable transcriptional changes were observed at 48 hours, with 21 genes overexpressed and 28 genes repressed in *Δbc1280*, including *tasA* and *calY* (log2FC values of -3.13 and -3.14, respectively) (Table S2). This result was further supported by RT‒qPCR analysis, which revealed lower expression levels for the genes located within the *eps1* and *eps2* genomic regions (Figure 2A). Our transcriptomic analysis additionally revealed that the expression of *bc2793* and *bc2794* pairs (log2FC values of -8.11 and - 8.43, respectively) was significant repressed. These uncharacterized loci are annotated as a putative Clp protease and an RNA polymerase ECF-type sigma factor, respectively. ECF sigma factors are known to transduce changes in the cell envelope into a bacterial genetic response^36,37^; thus, we hypothesized that under normal conditions, *bc2793* and *bc2794* positively regulate the gene expression of the main ECM components. The overexpression of *bc2793* and *bc2794* in *Δbc1280* and induction with a 10 μM solution of isopropyl β-D-thiogalactopyranoside (IPTG) significantly increased the biofilm biomass (Figure S2A). Considering that the putative protease (*bc2793*) could regulate the activity of *bc2794*, we overexpressed only the ECF factor, resulting in a six-fold increase in crystal violet staining at 100 µM IPTG compared to the strain without IPTG induction (Figure S2A). In contrast to these findings, the deletion of *bc2794* did not significantly affect biofilm formation, whereas the deletion of *bc2793*-*bc2794* resulted in a slight reduction in biofilm production compared to the wild-type strain, though this reduction was minimal relative to the biofilm-impaired phenotype observed in *Δbc1280* (Figure S2A). RT‒qPCR analysis revealed no significant changes in the expression levels of *tasA* or *calY* in *Δbc1280-*overexpressing *bc2794* or *bc2793*-*bc2794* compared to those in *Δbc1280* (Figure 2B). However, the relative expression levels of *eps1* and *eps2* were significantly upregulated in response to both genetic combinations. Thus, the ECF factor and its anti-sigma factor were associated with the regulation of exopolysaccharide expression in *B. cereus*, rather than the biofilm-related genes *tasA* and *calY*.

**Figure 2.**
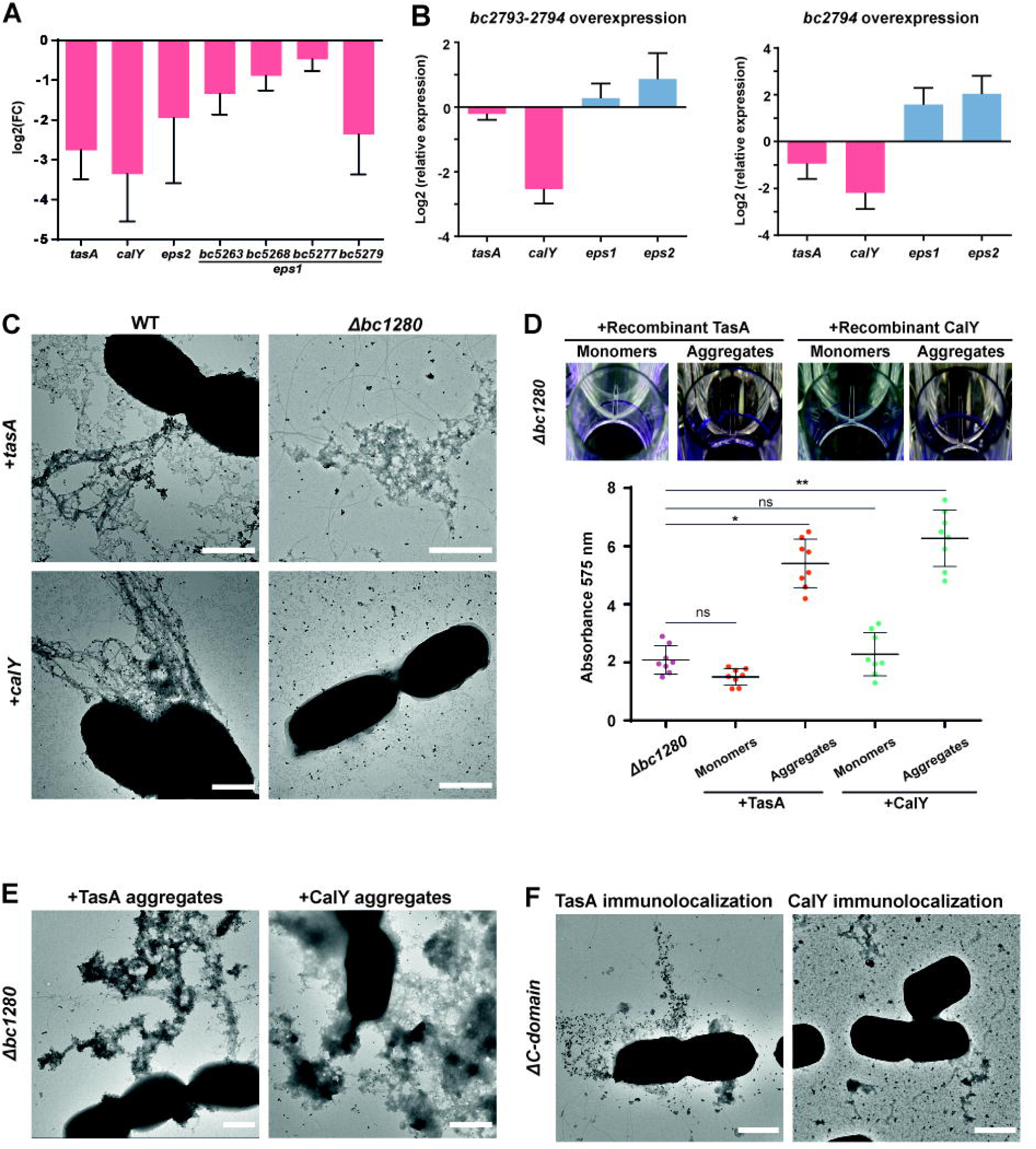
The biofilm-defective phenotype of *bc1280* is independent of the expression level of *tasA* or *calY.* **A)** Relative expression levels of *tasA*, *calY*, *eps2* and genes belonging to the *eps1* operon (*bc5263*, *bc5268*, *bc5277* and *bc5279*) obtained by RT‒qPCR in *Δbc1280* planktonic cells compared to wild-type cells at 48 hours. The average values of three biological replicates are shown with error bars indicating the standard deviation. **B)** Relative expression levels of *tasA*, *calY*, *eps1*, and *eps2* were measured in planktonic cells at 72 hours in the *Δbc1280* strain that overexpressed *bc2793-bc2794* or *bc2794* and compared to the wild-type strain. Average values from three biological replicates are presented, with error bars representing the standard deviation. **C)** Transmission electron micrographs of the wild-type and *Δbc1280* strains overexpressing *tasA* or *calY*, immunolabeled with specific anti-TasA or anti-CalY antibodies, respectively. Scale bar: 1 μm **D)** Extracellular complementation of *Δbc1280* after the addition of TasA or CalY at 6 μM in monomeric or fibrillar form. The biofilm mass was quantified by crystal violet staining, followed by absorbance measurement at 575 nm and comparison with the *Δbc1280* strain. Statistical analysis was performed using one-way ANOVA with Dunn’s multiple comparisons test. Statistical significance is indicated as p < 0.05 (*) and p < 0.01 (**). **E)** Phenotype reversion of *Δbc1280* upon the addition of 6 μM polymerized TasA or CalY, visualized by immunolabeling with anti-TasA and anti-CalY antibodies, respectively. Scale bar: 1 μm **F)** Negative-stained micrographs of TasA and CalY immunodetection in the *ΔC-domain* strain. Scale bar: 1 μm

In accordance with our hypothesis on the structural role of BC1280 in the polymerization of TasA or CalY, no reversion of phenotypes was observed in *Δbc1280* cells overexpressing *tasA, calY*, or both loci (Figure S2B). TEM analysis of negatively stained samples and immunolabeled with anti-TasA or anti-CalY antibodies revealed (Figure 2C and D): i) the presence of filaments from both proteins on the surfaces of WT cells, ii) aggregates reactive to anti-TasA antibodies in *Δbc1280* cells overexpressing *tasA,* and iii) the absence of filaments in *Δbc1280* cells overexpressing *calY* but with an extracellular signal associated with the anti-CalY antibody. We next examined whether TasA or CalY, which were produced by heterologous expression in *E. coli* and added exogenously to *Δbc1280*, might rescue the wild-type biofilm phenotype. The monomeric conformation of TasA or CalY failed to rescue biofilm formation; however, ring biomass formation was only partially restored when the proteins were incubated in vitro for one week to promote filaments assembly and then added at a final concentration of 6 µM (Fig 2D). In line with these findings, TEM analysis revealed that upon addition of TasA and CalY in their already assembled conformation, the cell surfaces were decorated with specific immunoreactive aggregates of these proteins (Figure 2E). Taken together, these data suggest that BC1280 is crucial for ECM maturation and raise the hypothesis that it might be involved in the polymerization of CalY or TasA. However, to fully understand the downregulation of *tasA* and *calY* over a longer period, additional research is required to ascertain whether this is a direct result of the absence of *bc1280* or an indirect consequence of an accumulation of TasA and CalY in the cytoplasm that are not being secreted.

Our previous findings indicated that the N-domain might partially fulfill the function of BC1280 in biofilm formation by specifically facilitating cell-to-cell interactions. Therefore, this domain may influence the final localization of TasA or CalY within the ECM. TEM analysis and immunolabeling of the *ΔC-domain* strain revealed that nanogold particles associated with TasA extracellular filaments were exposed on the cell surfaces; nevertheless, the morphologies of these filaments were different from those formed in the wild-type cells (Figure 2C and F). Nanogold particles corresponding to CalY monomers or oligomers, but not filaments, were observed in the extracellular matrix of the *ΔC-domain* strain (Figure 2C and F). Based on the alterations observed in the polymerization of TasA and especially CalY in the *ΔC-domain* strain, the C-domain of BC1280 might be involved in the polymerization of these proteins or their initial association with the cell surface.

### BC1280 is a cell wall-associated protein that partially colocalizes with CalY in the ECM

To precisely determine how BC1280 contributes to the assembly of the ECM, we further studied the localization of the protein at the cellular level. Our preliminary data indicated that BC1280 is likely expressed at very low levels and is undetectable by immunocytochemistry assays. Therefore, we conducted these experiments using a derivative *Δbc1280* strain that expressed a version of BC1280 fused with a histidine tag at the carboxyl terminal domain. The biofilm formation of *Δbc1280* was rescued by plasmid leakage without IPTG induction or upon induction with a 10 µM IPTG solution (Figure S3B). Furthermore, fractionation and immunodetection experiments with an anti-histidine antibody confirmed the presence of BC1280 in the extracellular matrix, cell wall, and cellular fractions (membrane and cytosolic content) (Figure 3A). TEM analysis and immunolabeling using anti-histidine antibodies revealed that foci of nanogold particles related to BC1280 were distributed regularly along filaments that emerged from the cell surfaces (Figure 3A). To better determine the localization of BC1280 in the cell wall, we conducted confocal laser scanning microscopy (CLSM) and immunocytochemistry using anti-histidine and secondary antibodies conjugated to Atto 488 and double staining with wheat germ agglutinin (WGA) and Hoechst to visualize the cell wall and DNA, respectively. According to the results of the fractionation experiments, a limited number of foci corresponding to BC1280 proteins fused to the histidine tag per cell were observed in association with the cell surface (Figure 3C and S3C). Considering the relevance of the N-domain to the functionality of BC1280, we also studied its contribution to the cellular localization of whole BC1280. Based on crystal violet staining of the biomass that had adhered to the well, the amount of N-domain expressed without IPTG induction or after induction with 10 μM IPTG solution was sufficient to rescue the biofilm phenotype of *Δbc1280* (Figure S3B). Cellular fractionation and immunoblotting assays demonstrated a specific reactive band in the extracellular matrix and cellular fractions and an undetectable signal associated with the cell wall (Figure 3B). In addition, visualization via TEM confirmed the presence of N-domain-related foci along the filaments emerging from the cells, as previously observed for the entire protein, but no detectable fluorescent signals associated with the cell wall were observed via CLSM (Figure 3B and C). The disparities noted between the full-length and the truncated form of the protein suggest that the C-domain contributes to the retention of BC1280 in the bacterial cell wall, likely before BC1280 reaches the ECM.

**Figure 3.**
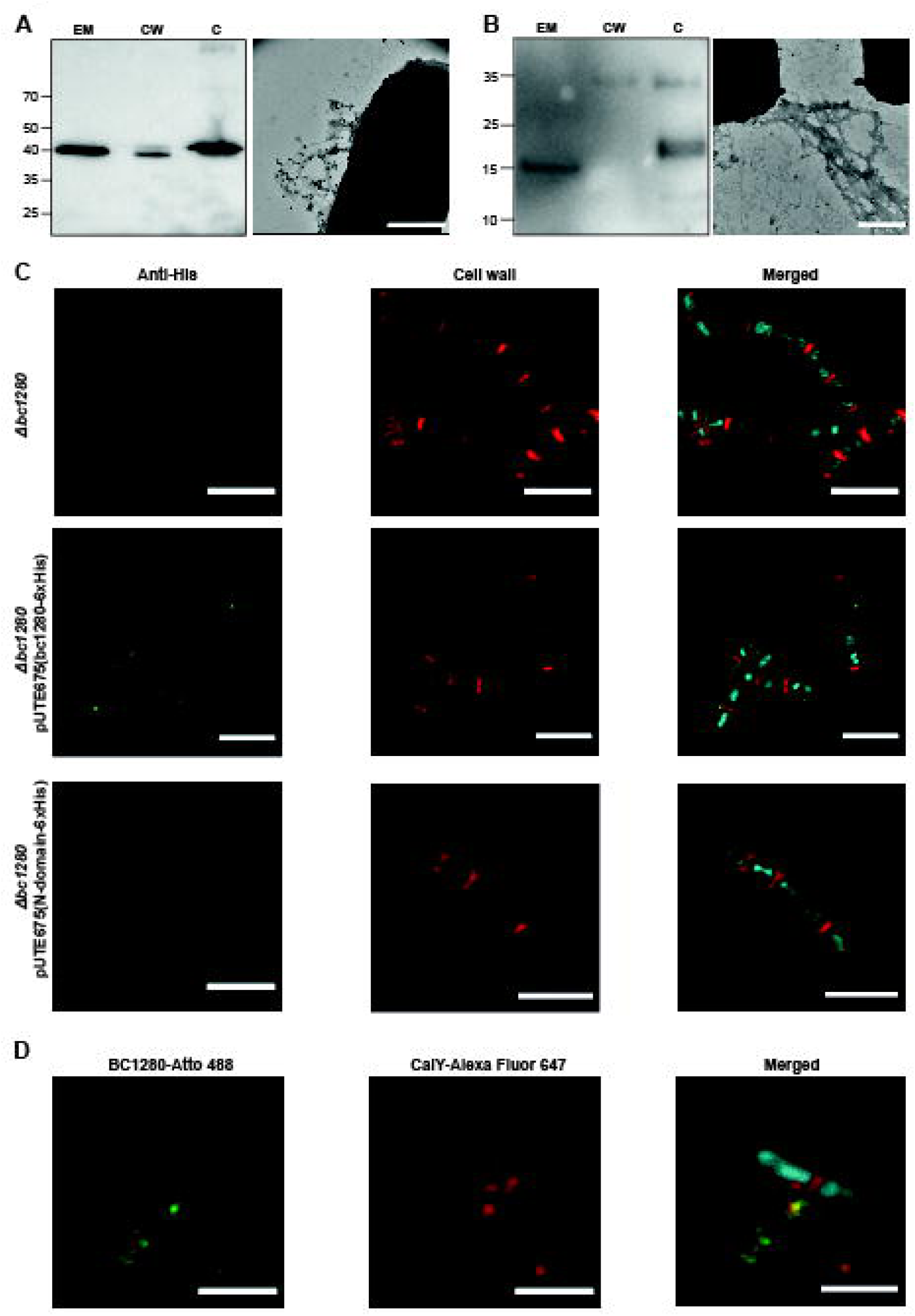
BC1280 is a cell wall-associated protein that partially colocalizes with CalY in the ECM. **A)** and **B)** Left: Immunodetection by Western blotting was performed for BC1280 (A) and the N-domain (B) fused to a histidine tag in different cellular fractions of *Δbc1280*. This assay was conducted under conditions of overexpression using the vector pUTE657 and with the addition of 10 μM IPTG. Biofilm samples were grown for 48 hours, after which the resulting biomass was recovered for fractionation into EM (extracellular medium), CW (cell wall), and C (membrane and cytosol) fractions. The immunoreactive bands were detected using an anti-His antibody (1:2500). The immunoblot images were edited by cropping and splicing for better illustration. Right: Negative stained images of BC1280 (A) and the N-domain (B) in the ECM of the *Δbc1280* strain. The proteins were immunolabeled using an anti-His antibody (1:100) as the primary antibody and a goat-antirabbit secondary antibody conjugated with 20 nm nanogolds (1:100). Scale bar: 500 nm **C)** Immunolocalization of BC1280 and the N-domain determined using CLSM. Biofilm samples were processed using immunochemistry, with anti-His antibody (1:100) serving as the primary antibody and a secondary antibody conjugated to Atto 488. The localization of BC1280 and the N-domain is shown in green. The cell wall is labeled in red using WGA, and the DNA is stained in blue using Hoechst. Scale bar: 5 μm **D)** Immunolocalization images of BC1280 and CalY obtained through CLMS. An anti-His antibody (1:100) and a secondary antibody conjugated with Atto 488 (1:400) were used for the immunolabeling of BC1280. A specific primary anti-CalY antibody (1:100) and a secondary antibody labeled with Alexa Fluor 647 (1:400) were used for CalY detection. Additionally, DNA was stained with Hoechst (1:1000) to visualize the bacteria, which are shown in blue. Scale bar: 3 μm

To further define the connection between BC1280 and CalY or TasA at the cellular level, we studied their colocalization via immunocytochemical analysis of cells expressing pUTE657-*bc1280* fused to a histidine tag using specific primary antibodies. CLMS analysis revealed that CalY and BC1280 colocalized along the fibrils that compose the ECM (Pearson’s coefficient of 0.58 and a p value of 0.0002) (Figures 3D and S3D). In most cases, the signal corresponding to CalY was homogeneously distributed, and BC1280 was detected as discrete foci dispersed along the filaments. However, no colocalization signal was detected for TasA or BC1280, with a Pearson’s coefficient of 0.19 and a p value of 0.0002 (Figure S3E and F). These findings were supported by Mander’s coefficient, which represents the percentage of overlapping pixels for each fluorescent channel. For the BC1280-CalY pair, approximately 65% of the pixels overlapped in both channels; for BC1280-TasA, the overlap decreased to 26% for Atto 488 and 21% for Alexa Fluor 647 (Figure S3F). The observation of BC1280 and CalY colocalizing within the ECM hints at a potential structural relationship, including possible direct interactions, between these proteins.

### Heterologously produced BC1280 oligomerizes into macromolecular complexes driven by the N-domain

BC1280 and the truncated version comprising the N-terminal domain (from A^39^ to G^190^) were purified by heterologous expression in *E. coli* Lemo21(DE3) and subjected to further structural analyses (Figure 4A). The two versions of the protein were purified under soluble conditions by affinity chromatography followed by size exclusion chromatography, and by tandem mass spectrometry analyses, the amino acid sequences of each protein were confirmed (Figure S4A). The two fractions collected at different elution volumes in the size exclusion chromatogram suggested that in buffer containing 20 mM Tris and 50 mM NaCl at pH 7, BC1280 and N-domain39-190 could assemble into two main molecular entities with distinct oligomeric states (Figure 4B and E). Dynamic light scattering (DLS) analysis showed that both protein assemblies exhibited a degree of polydispersity, with particles ranging in diameter from ∼10 to 600 nm (Figure 4C-D and F-G for BC1280 and N-domain39-190, respectively; for further details, see Table S3). Remarkable differences were observed in the second elution sample; specifically, the N-domain39-190 exhibited a greater proportion (∼53%) of small oligomers than did BC1280 (∼18%) (Figure 4D and G). To complement the DLS results, each fraction was resolved in a native gel (Figure 4H). The sample corresponding to fraction 1 of each protein, which presumably contained larger aggregates, failed to enter the gel, suggesting that the molecular complex was larger than 1000 kDa. The second elution fraction was resolved as a band at 480 or 240 kDa for BC1280 or N-domain39-190, respectively, which corresponds to oligomers of approximately 15 subunits each. TEM analysis confirmed that fraction 1 contained large aggregates of varying sizes for both proteins (Figure S4B). The samples corresponding to the second elution fraction contained macromolecular complexes that were similar but smaller in diameter (Figure S4B). Taken together, these results indicate that the variability in the oligomerization of the full-length protein is preserved in the truncated version that contains only the N-terminal region.

**Figure 4.**
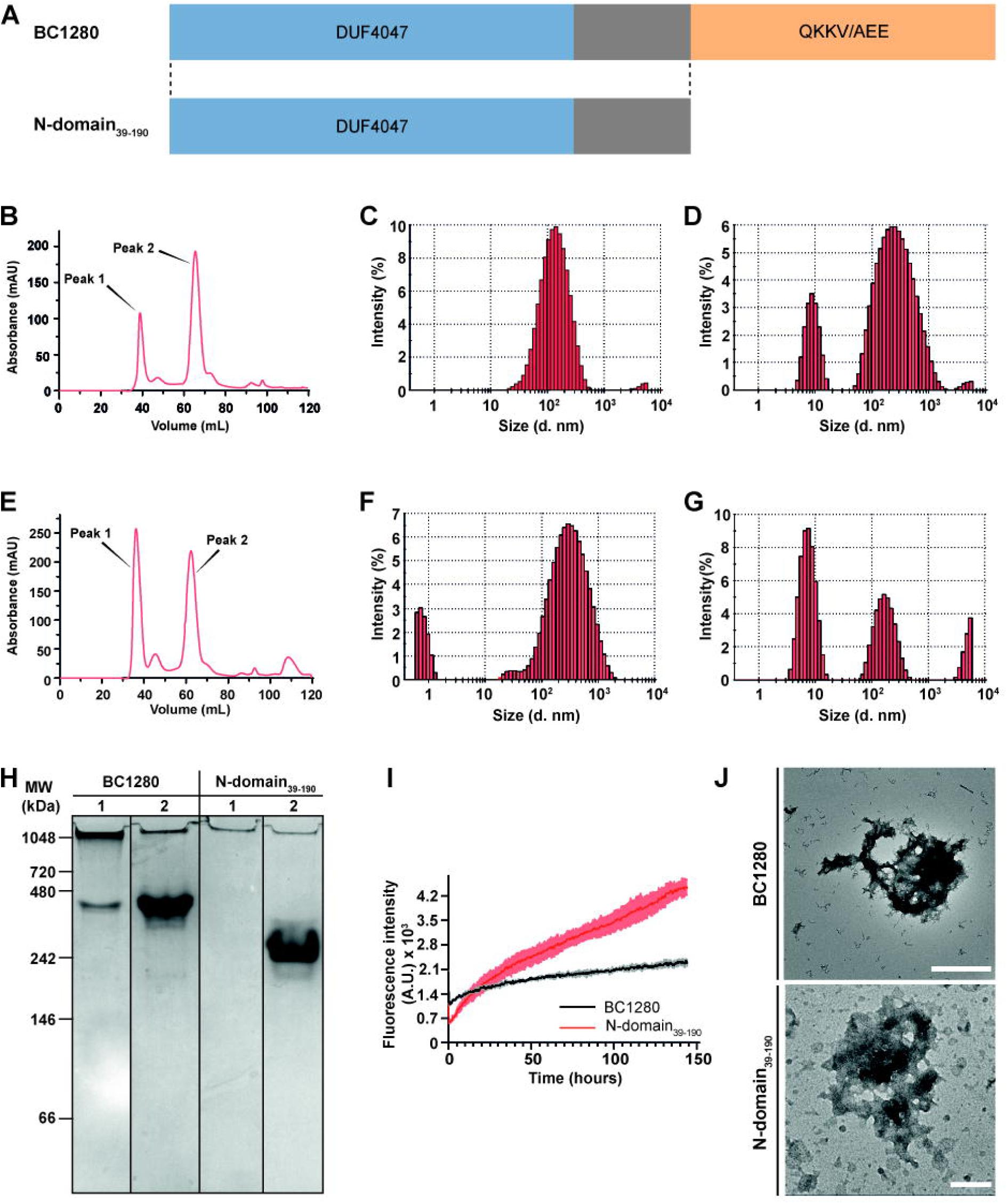
Heterologously produced BC1280 oligomerizes into macromolecular complexes driven by the N-domain. **A)** Scheme illustrating the two constructs based on the full-length sequence of BC1280. Each construct was fused to a poly-histidine tag at the C-terminal domain, and the resulting two recombinant proteins were produced through recombinant expression in *E. coli*. **B)** Size-exclusion chromatogram obtained after BC1280 was purified under native conditions. **C)** and **D)** Size distribution histograms obtained via DLS showing the diameters of the aggregates formed by BC1280 for peaks 1 and 2, respectively, as determined by SEC. **E)** Size-exclusion chromatogram obtained after purifying N-domain39-190 under native conditions. **F)** and **G)** Size distribution histograms obtained via DLS, illustrating the diameters of the aggregates formed by N-domain39-190 for peaks 1 and 2, respectively, as determined by SEC. **H)** Native polyacrylamide gel showing the oligomers formed by BC1280 and N-domain39-190 following native purification through heterologous expression in *E. coli*. The numbers on each line correspond to the first and second peaks, respectively, that were detected in the size-exclusion chromatograms of each sample. The gel lanes were cropped and spliced for illustrative purposes. All the lines superimposed on the image delineate the lanes and boundaries of the spliced images. **I)** Aggregation kinetics of BC1280 (black curve) and N-domain39-190 (red curve) determined using ThT fluorescence. **J)** Transmission electron micrographs of BC1280 and N-domain39-190 at the end of the polymerization experiment shown in I. Scale bar: 200 nm

To further delineate the dynamics of oligomerization, we focused on the second fraction, which was composed mainly of small oligomers and could promote the formation of larger molecular entities. The fluorescence signal emitted upon binding to thioflavin T (ThT)^38^ showed a fast polymerization curve without a lag phase, aligning with the presence of oligomers under native conditions (Figure 4I). Interestingly, the slope of the exponential phase was more pronounced for the N-domain39-190, and the fluorescence intensity signal was twofold greater than that of the full-length protein after 150 hours of incubation. These results correspond with the higher percentage of small oligomers formed by the N-domain39-190 (via DLS analysis), which suggested that the truncated protein seeds rapidly. Furthermore, TEM analysis of negatively stained samples revealed macroscopic particles of variable sizes and an amorphous aggregated morphology for both proteins, resembling the aggregates observed for CalY but not the fibrillar structures formed by TasA (Figure 4J).^32^ These results strongly indicate that the N-domain is involved in protein oligomerization and suggest that the absence of the C-domain in the truncated protein might facilitate a faster aggregation rate.

### The assembled N-domain39-190 contains a rigid core with two β-strand-rich regions, while the C-domain increases global rigidity

The amyloid-like nature of TasA and CalY^32^, when heterologously produced in *E.coli*, along with the association of BC1280 with these two proteins *in vivo* and its tendency to oligomerize *in vitro*, led us to suspect an amyloid-like nature for BC1280. Since the N-domain retains the functionality of BC1280 and the ability to form oligomers, this version of the protein was considered for our initial structural studies. The X-ray diffraction pattern of the centrifuged pellet of the N-domain39-190 assemblies displayed typical features reminiscent of the cross-β structure of amyloids, with strong signals at 4.7 Å and a slight signal at ∼10-11 Å, corresponding to the inter-strand and inter-sheet spacings, respectively^39^ (Figure 5A). In light of these findings, we predicted protein aggregation and amyloidogenicity based on the amino acid sequence of BC1280 using the bioinformatics tools AmylPred2,^40^ FoldAmyloid,^41^ MetAmyl,^42^ PASTA 2.0^43^ and TANGO.^44^ At least two algorithms predicted the following amino acid stretches as putative aggregative regions and amyloid-forming segments: i) L^51^STAIIFP^58^, ii) I^98^S^99^, iii) L^115^QNVYT^120^ and iv) V^144^SITVN^149^ (Figure S4C; letters in bold). Based on bioinformatics analysis using FoldUnfold,^45^ the N-domain appears intrinsically folded but the C-domain is predicted to be unstructured, possibly due to the presence of repetitive amino acid sequences rich in Gln, Lys, and Glu, which are disorder-promoter residues^35,46^ (Figure S4C). As the propensity to form β-sheet-rich self-assembled structures is often associated with regions enriched in hydrophobic residues, we analyzed the hydrophobicity pattern of BC1280 using ProtScale.^47^ The peptide T^50^LSTAIIFPKT^60^, previously identified as a putative amyloidogenic segment, was found to be highly hydrophobic (Figure S4D). Finally, AlphaFold^48,49^ was used to obtain a tentative secondary structure. The model predominantly exhibits an α-helix conformation with low confidence value, except for a small central helical sandwich. This contrasts with the enrichment in β-sheet structural elements derived from ThT polymerization kinetics and the cross-β structure observed by X-ray diffraction (Figure S4E). We further performed fine structural characterization of the molecular conformation of these protein assemblies using magic-angle spinning (MAS) SSNMR, which has previously been employed to study non-crystalline self-assemblies, such as amyloids.^50–52^ Uniform ^13^C/^15^N-labeled N-domain39-190 assemblies were produced to improve NMR sensitivity and perform multidimensional MAS SSNMR studies. To investigate the presence of a central rigid core within the assemblies, we used dipolar-based polarization transfer (cross-polarization) and two-dimensional ^13^C-^13^C spectroscopy. A 50 ms mixing time proton-driven spin-diffusion (PDSD) experiment was used to detect intra-residue ^13^C-^13^C correlations from the rigid core of the assembly. For the spectral analysis, we first attempted to exploit the secondary structure model predicted by AlphaFold,^48,49^ and the Cα and Cβ chemical shifts were predicted using SPARTA.^53^ Subsequently, the predicted spectrum was superimposed with the experimental 2D ^13^C-^13^C spectrum acquired for N-domain39-190 (Figure S5A). No correlation was found between the experimental and predicted NMR spectral patterns, which reflects the discrepancies between the AlphaFold model and the structure of the experimentally obtained multimers. These findings highlight that, while bioinformatics predictions might reflect the relevant structures under certain conditions, differences between predicted and experimentally observed structures can result from factors such as the nature of the sample, the buffer composition (pH or salt), or oligomerization. The NMR data provide evidence for the propensity of BC1280 to undergo structural changes upon assembly under certain *in vitro* or environmental conditions. Thus, we manually inspected the SSNMR cross-peaks and identified ∼35-40 spin systems (i.e., corresponding to ∼35-40 residues), as exemplified for an isoleucine in Figure 5B. Following well-established statistical data for Cα and Cβ chemical shifts according to their secondary structure,^54^ we noticed that these residues are involved mainly in β-sheet secondary structure elements. This finding was illustrated by the isoleucine spectral pattern observed with Cα and Cβ chemical shift values of ∼60 and 42 ppm, respectively, which are indicative of a β-sheet conformation (Figure 5B). Approximately 35-40 out of the 151 amino acids comprising the N-domain39-190 were detected within the rigid core by SSNMR, suggesting that the rigid core encompasses only a portion of the N-domain39-190 sequence. However, the presence of residues undetectable in dipolar-based SSNMR experiments is not surprising, as this observation has been made for most pathological and functional amyloids and self-assemblies to date.^55,56^

**Figure 5.**
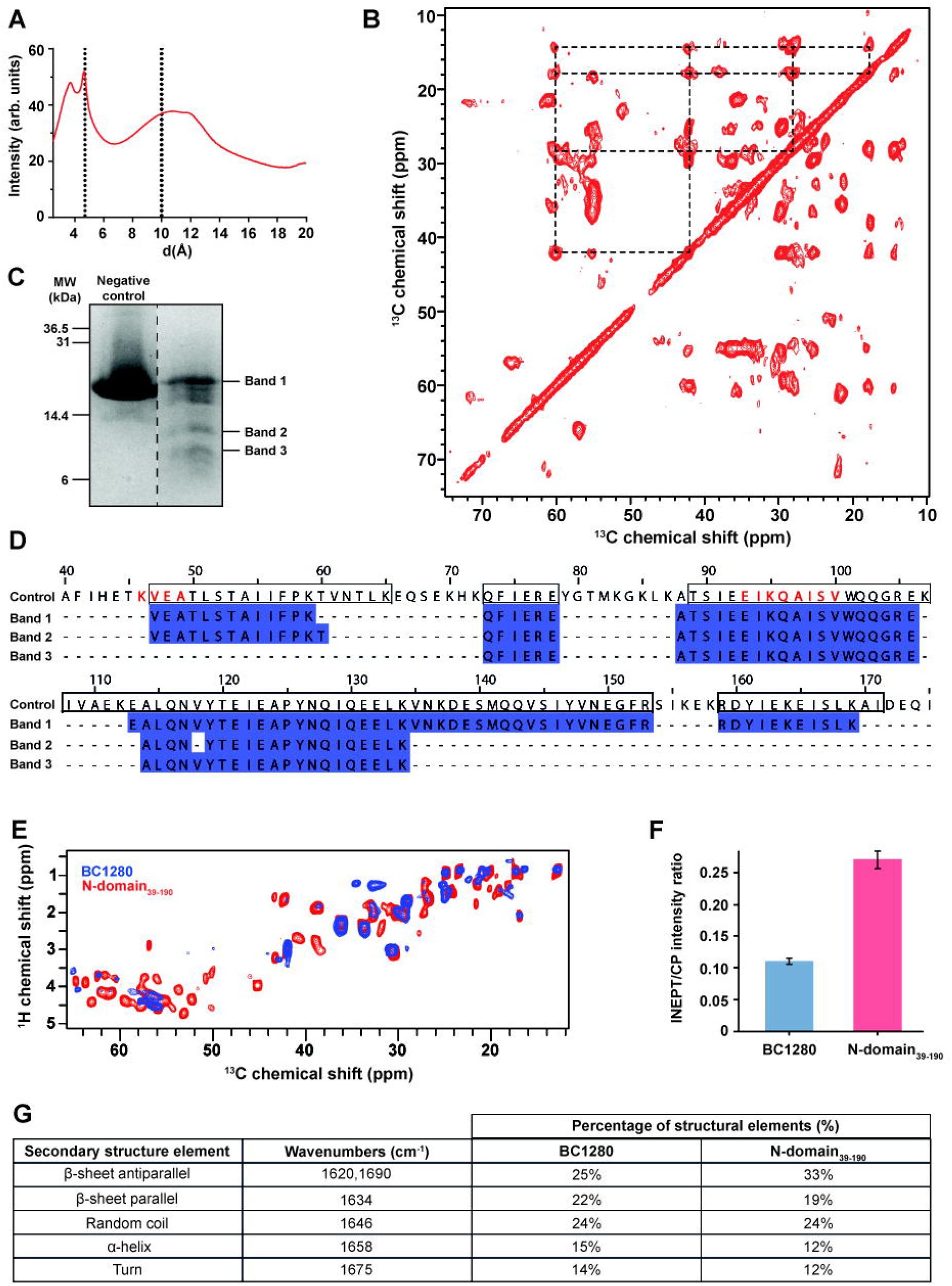
The assembled N-domain39-190 contains a rigid core with two β-strand-rich regions, while the C-domain increases global rigidity. **A)** X-ray diffraction signals of N-domain39-190 at 4.7 and 10 Å, indicating interstrand and intersheet spacing, respectively. A) 2D ^13^C-^13^C PDSD experiments at 50 ms for the N-domain39-190. The chemical shifts that correspond to isoleucine residues in the β-sheet conformation are highlighted. **C)** A 12% SDS‒PAGE gel was obtained after the treatment of N-domain39-190 aggregates with proteinase K for 45 minutes. The three bands marked were analyzed by tandem mass spectrometry. **D)** Determination of the rigid core of N-domain39-190 by tandem mass spectrometry. MS data were obtained from the bands selected after digestion of N-domain39-190 with proteinase K. The amino acids outlined by a box are the rigid regions identified in the absence of proteinase-K, and letters in red correspond to the residues identified by SSNMR PDSD at 50 and 200 ms. **E)** Superimposition of the ^1^H-^13^C INEPT spectra of BC1280 (blue) and N-domain39-190 (red), recorded at a 1H frequency of 600 MHz (276 K and 96 scans for BC1280; 300 K and 640 scans for N-domain39-190) showing signals corresponding to the flexible part. **F)** Estimation of the global rigidity based on the INEPT:CP ratio of chemical shifts for each sample. **G)** Secondary structure content of the BC1280 and N-domain39-190 aggregates probed via ATR-FTIR analysis.

In light of these findings, we observed that the spectral resolution, as measured by the ^13^C line width, is ∼60-120 Hz (full width at half-height), which indicates a relatively well-ordered of the subunits within the complex. These observations are consistent with our previously reported ^13^C line widths for *B. subtilis* TasA (∼30-100 Hz), *B. subtilis* TapA (∼200 Hz) and *B. cereus* CalY (∼200 Hz), which, respectively form well-ordered fibrils (TasA) and polymorphic aggregates (TapA and CalY) in vitro.^32^ To gain more site-specific insights into the rigid core of the N-domain39-190, the protein was incubated with proteinase K, the bands corresponding to the rigid regions were resolved via SDS‒PAGE, and the amino acid sequences were determined through tandem mass spectrometry analysis (Figure 5C and D). Interestingly, proteinase K digestion suggested that the following resistant regions were present: the highly hydrophobic region V^47^-K^59^, which contains the Ala-Ile-Ile segment, and the region A^88^-K^134^. Further examination of the SSNMR data using an additional 200 ms PDSD experiment (Figure S5C) led us to identify several rigid protein segments. Specifically, these segments correspond to the regions identified through proteinase K digestion analysis, namely, K^46^VEA^49^ and E^93^IKQAISV^100^. Notably, the total number of residues identified as resistant to proteinase K was slightly greater than the number of spin systems identified by SSNMR. This difference suggests that the core visible in SSNMR contains fewer residues compared to the regions identified as proteinase K resistant. Alternatively, this observation may result from insufficient spectral resolution to detect all correlations. Based on these results, it can be inferred that the N-domain39-190 contains a structured core with a significant presence of rigid residues in a β-sheet conformation.

As the full-length protein exhibits distinct polymerization kinetics from those of the N-domain39-190, we conducted a comprehensive biophysical characterization of full-length BC1280 to identify putative differences from the truncated N-domain39-190 in the context of macromolecular assemblies. The full-length protein sample underwent a similar treatment as the N-domain39-190 sample, and upon ultracentrifugation, the following notable differences were observed: i) BC1280 assembled into a transparent gel-like pellet with high viscosity, while ii) the N-domain39-190 pellet aggregated into a non-viscous white precipitate. Consistent with previous findings for the N-domain39-190, the diffraction pattern of the BC1280 assemblies showed an intense signal at ∼4.7 Å and a less pronounced peak at ∼10 Å (Figure S5D). The C-domain of BC1280 consists of fifteen tandem repeats that were predicted to be disordered; thus, we wondered whether this domain participates in the overall folding of the full-length protein assemblies. BC1280 was uniformly ^13^C labeled, and the resulting aggregates were analyzed via MAS SSNMR. To compare the molecular conformation of N-domain39-190 to that of full-length protein assemblies, we superimposed 2D ^13^C-^13^C SSNMR spectral fingerprints and observed significant overlap, suggesting that the rigid core of the N-domain is highly conserved in full-length protein assemblies (Figure S5E). As only a few additional NMR signals were observed in the full-length protein sample, we hypothesize that the C-domain is not a primary component of the rigid structural fold of the BC1280 assemblies.

To complement the SSNMR analysis of the rigid BC1280 core, we used dynamics-based spectral editing via J-coupling polarization transfer to detect dynamic protein segments^57^ and gain insights into any dissimilarities between the two protein versions. The superimposition of 2D ^1^H-^13^C-insensitive nuclei was enhanced by polarization transfer (INEPT) of the BC1280 and N-domain39-190 assemblies, and the results revealed a significant difference in the spectral fingerprint between the two samples (Figure 5E). More INEPT signals were observed in the N-domain39-190 than the full-length protein, suggesting that the absence of tandem repetitions in the C-domain results in a more flexible conformation overall. To confirm this hypothesis, we estimated the degree of rigidity by comparing signal intensities from 1D ^13^C-detected INEPT and CP experiments (Figure 5F), respectively, by probing mobile and rigid protein segments through the INEPT/CP intensity ratio. We observed a different behavior, with a decrease in mobile signals in BC1280 (ratio ∼0.1) compared to N-domain39-190 (ratio ∼0.25), suggesting fewer dynamic regions in the assemblies of the full-length protein.

Considering the differences in the overall rigidity between BC1280 and N-domain39-190, we estimated the relative amount of secondary structural elements using attenuated total reflection-Fourier transform infrared spectroscopy (ATR-FTIR) analysis. To identify the minimum peaks within the amide I range according to well-established assignments,^58^ the second derivative was calculated from the raw data, and deconvolution was performed to determine the percentage of contribution for each secondary structure type (Figure 5G and S5F, G and H).Overall, due to the additional structural elements, the ATR-FTIR spectra of BC1280 was more challenging to fit. The N-domain39-190 exhibited 33% antiparallel β-sheets, while the full-length protein had 25% antiparallel β-sheets and a slightly greater proportion of parallel β-sheets. Both proteins displayed an equal overall percentage of secondary structure in the random coil conformation, while BC1280 contained slightly more α-helices and turns than the N-domain39-190.

### BC1280 acts as a chaperone for CalY, with its functionality modulated by the C-domain

The observed in vivo colocalization of BC1280-CalY in the extracellular matrix and the potential involvement of BC1280 in CalY polymerization led us to ask whether BC1280 might act as an accessory protein that influences CalY assembly. We examined how BC1280 affects CalY by studying the aggregation kinetics of CalY at different BC1280 molar ratios using ThT fluorescence emission. Surprisingly, we observed that even at ratios (CalY:BC1280) of 70:30 or 50:50, the polymerization kinetics were significantly affected compared to CalY incubated alone (Figure 6A). This negative impact, together with the fact that BC1280 expression is likely tightly regulated, prompted us to investigate the relative levels of BC1280 and CalY during biofilm formation using an isobaric tag for relative and absolute quantification (iTRAQ) analysis (Figure 6B). At 24 hours, the relative accumulation rate of CalY was approximately 3.3 times higher than that of BC1280; nevertheless, BC1280 accumulated by 48 hours, reaching a ratio (CalY:BC1280) of 2:1. This information was used to re-estimate the protein ratios for copolymerization assays of CalY with BC1280. The presence of low concentrations of BC1280 (2 and 10 μM) did not impact the polymerization kinetics of a 40 μM solution of CalY (Figure 6C). In contrast, the presence of only the N-domain39-190 significantly reduced the rate of polymerization kinetics of CalY, resulting in a lower fluorescence intensity at the end of the experiment compared to CalY alone (Figure 6C). In addition, the negative impact on CalY polymerization was amplified in the presence of non-freshly purified full-length protein or N-domain39-190, which are enriched in preformed oligomers (Figure S6B and C). The samples corresponding to CalY incubated with a freshly purified 10 µM solution of BC1280 or N-domain39-190 were visualized via TEM. Consistent with the ThT results, incubation of 10 μM BC1280 with CalY resulted in macromolecular aggregates similar to those formed by CalY (Figure 6D). However, the incubation of CalY with the N-domain39-190 led to the accumulation of small oligomers (Figure 6D), aligning with the previous *in vivo* results where the absence of the C-domain (*ΔC-domain*) arrested the formation of CalY filaments (Figure 2F). The ability of BC1280 to regulate the CalY polymerization rate points to a potential chaperone role for BC1280 in a concentration-dependent manner. This hypothesis was further supported by circular dichroism analysis, which revealed significant structural changes in CalY in the presence of BC1280 in a concentration-dependent manner. After 16 hours of incubation under agitation, a 10 μM solution of BC1280 induced an increase in the β-sheet conformation of CalY compared to CalY alone or after incubation with 2 μM BC1280, which predominantly appeared as a random coil or a disordered conformation (Figure 6E).

**Figure 6.**
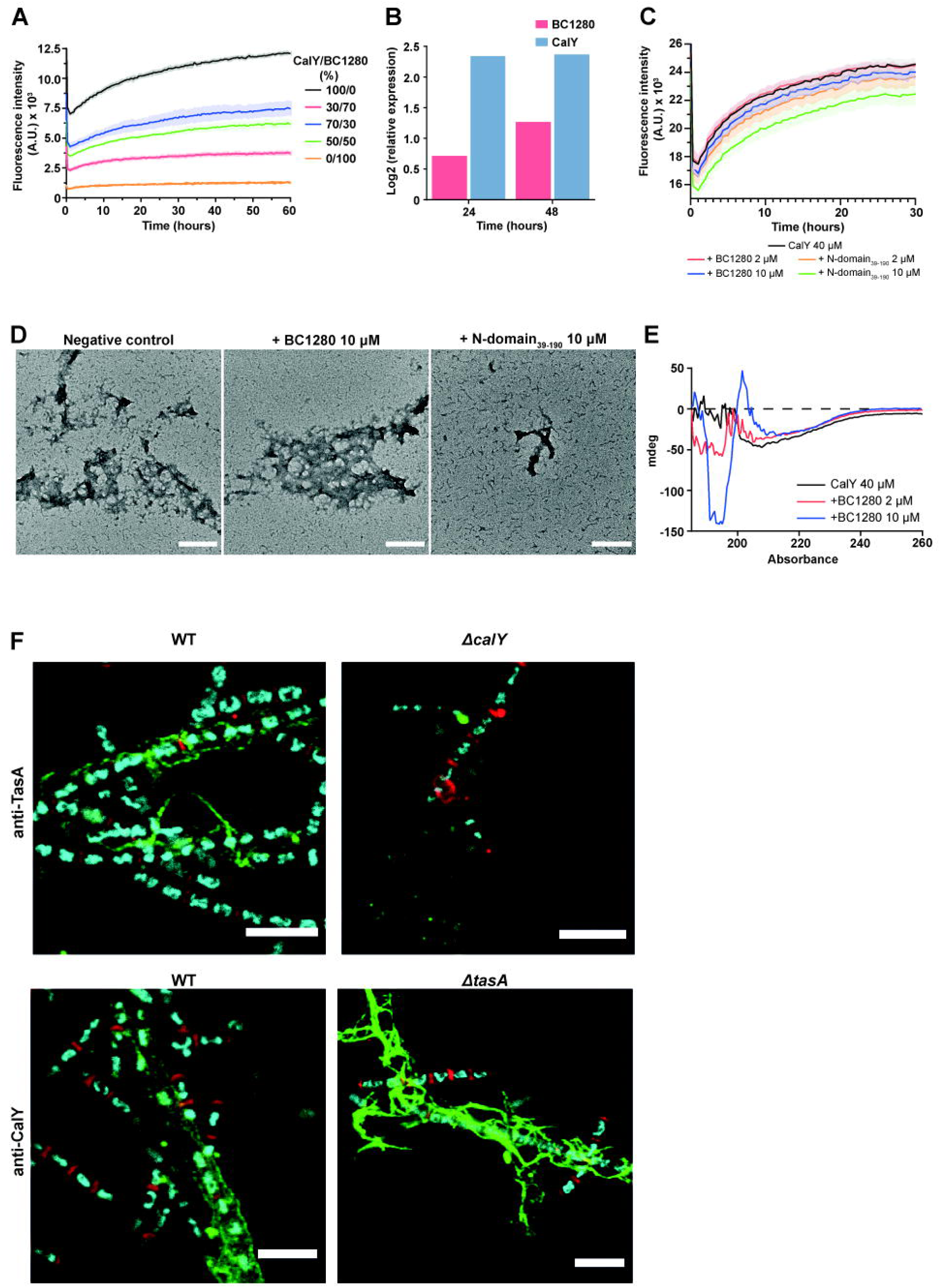
BC1280 acts as a chaperone for CalY, with its functionality modulated by the C-domain. **A)** Polymerization kinetics of different CalY/BC1280 molar ratios measured by ThT fluorescence emission. Error bars represent the SEM. **B)** Relative levels of BC1280 and CalY in biofilm cells during biofilm formation using iTRAQ. **C)** Kinetics of ThT binding of CalY in the presence of BC1280 or the N-domain39-190 at concentrations of 2 and 10 µM. Error bars represent the SEM. **D)** Transmission electron micrographs of negatively stained samples of CalY aggregates in the presence of BC1280 or N-domain39-190 at a concentration of 10 µM after 30 hours of incubation at 37°C. Scale bar: 200 nm **E)** Circular dichroism (CD) spectra of CalY incubated for 16 hours under agitation alone or in the presence of BC1280 at 2 and 10 μM. **F)** Immunolocalization of TasA and CalY in wild-type and mutant strains using CLSM with specific primary antibodies against TasA and CalY, and a secondary antibody conjugated to Atto 488. Green fluorescence represents TasA and CalY localization, with the cell wall stained red (WGA) and DNA stained blue (Hoescht). Scale bar: 5 μm

The experimental data shown in this work strongly suggest a direct interaction between BC1280 and CalY, yet the contribution of TasA within the filaments that support the ECM of *B. cereus* remains uncertain. In vitro co-assembly experiments have demonstrated that CalY can function as an accessory protein in the assembly of TasA, without altering its structural arrangement.^32^ Moreover, previous findings from our laboratory indicated that deletions in *tasA* or *calY*, while containing the *bc1280* gene, did not completely arrest biofilm formation; instead, mutant strains displayed biofilms with distinct phenotypes.^25^ Therefore, we addressed the distribution of their respective paralogue protein within the ECM in *ΔtasA* and *ΔcalY* mutants compared with the wild-type strain. Immunochemistry and CLMS analysis revealed fibrillary structures reactive to anti-TasA antibodies decorating the surfaces of wild-type cells, whereas discrete foci of TasA fluorescence signals were observed in the ECM of *ΔcalY* (Figure 6F). Remarkably, in *ΔtasA* mutant strain, CalY protein exhibited a higher degree of fibrillary organization compared to the immunofluorescence signal observed in *B. cereus* wild-type cells (Figure 6F). These results provide evidence that CalY can polymerize into filaments in *ΔtasA* in the presence of BC1280. In contrast, its paralogue TasA doesn’t polymerize into filaments *in vivo* when CalY is absent, highlighting the high complexity of the mechanism underlying the ECM assembly in the human pathogen *B. cereus*.

## DISCUSSION

Bacterial extracellular filaments are fascinating polymers from structural and biological perspectives. The diversity of structures, sizes, functions and mechanisms of polymerization reflects the specificity of their functions and need for tight regulation. In this work, we pinpointed BC1280 as a specific biofilm orchestrator factor that is exclusive to the *B. cereus* group. In addition, we propose an intriguing molecular mechanism by which this protein could contribute to the assembly of TasA-CalY filaments on the cell surface of this opportunistic human pathogen. The severe defective phenotype observed in *Δbc1280*, when contrasted with the phenotypes of *ΔtasA* and *ΔcalY*, substantiates the indispensable role of BC1280 in biofilm maturation.^25^

While BC1280 could be compared with the chaperone TapA from the phylogenetically related strain *B. subtilis*, notable differences between the two proteins highlight their distinct characteristics. First, BC1280 is exclusive to biofilm cells, whereas TapA is expressed in the planktonic population and from the early stages of biofilm formation. Upon *tapA* deletion, altered and fewer filaments are produced due to a drastic decrease in TasA levels, while the transcriptional expression of *tasA* remains unaffected.^59^ Remarkably, our findings reveal that *tasA* and *calY* levels are transcriptionally downregulated in *Δbc1280*, resulting in the absence of filaments. It is important to highlight that in a previous study by our group, we observed that the heterologous expression of *B. cereus* alleles in the *B. subtilis ΔtapA* strain only reverts biopellicle formation when *sipW-to-calY*, including *bc1280*, are overexpressed, indicating only partial functional and mechanistic overlap of TapA and the proteins encoded in *sipW- to-calY*.^25^ Moreover, in the same study, it was demonstrated that in the *B. subtilis ΔtasA* mutant, in which *tapA* is still expressed, when only *sipW-tasA* or *sipW-calY* from *B. cereus* are overexpressed without the *bc1280* allele, the biopellicle is not formed, clearly evidencing that the mechanism of assembly of TasA and CalY is specific of BC1280 and cannot be replaced by TapA from *B. subtilis*.^25^ The idea of a partial mechanistic overlap between BC1280 and TapA is further corroborated by the structural differences observed in the predicted model of BC1280 and the solved structure of TapA.^60^ While considering that structures exhibit context-dependent variations, they were partially predicted with high confidence, supporting the notion that these structures should occur in a specific in vivo configuration and perform functions such as membrane- or cell-wall interactions.

The results of this study demonstrate that BC1280 contains distinct structural and functional regions: the N-terminal half, which is sufficient to fulfill the functionality of BC1280; and the C-terminal half, which appears to be involved in attachment to the cell surface and also controls the interaction of BC1280 with CalY. After establishing that BC1280 is an ECM protein and its association with the cell surface, along with its functional relationship with CalY, we propose its role as a chaperone for CalY based on the following lines of evidence: i) the biofilm phenotype remains unchanged even after the overexpression of *tasA* and/or *calY* in the *Δbc1280* mutant strain; ii) filaments of CalY are not formed in the *ΔC-domain* strain; iii) colocalization of BC1280 and CalY within the ECM occurs *in vivo;* and iv) BC1280 induces a structural rearrangement of CalY in vitro. Collectively, these findings prompted us to rename BC1280 as CapP, a CalY-assisting polymerization protein. In addition to its fundamental role in controlling pilus assembly, CapP functions as a checkpoint of biofilm maturation by regulating the expression of EPS via an uncharacterized pair of Clp-proteases and cognate ECF-type sigma factors. However, this is a novel and specific biofilm regulatory pathway that deserves further investigation.

We determined in vitro that the N-domain of CapP might trigger or support protein aggregation, leading to the formation of a rigid core of ∼ 40 amino acids enriched in β-sheets. Through SSNMR, we determined at the atomic level that the rigid core contains at least the following rigid regions: K^46^VEA^49^ and E^93I^KQAISV^100^. The role of each amino acid stretch remains to be elucidated; nevertheless, we propose that the K^46^VEATLSTAIIFP^58^ region significantly influences the aggregation propensity of CapP in vitro, considering that i) the sequence is situated in the most hydrophobic region of the protein, ii) the residues within this region are highly conserved, and iii) the region was detected in the proteinase K-resistant fraction. Although the core is conserved in the full-length protein, it exhibits slower polymerization kinetics and a greater polydispersity index than that of the truncated N-domain. In addition, a higher global rigidity is generated by the C-domain in CapP compared to the truncated N-domain. Together, these findings strongly suggest that the entire protein has a more complex folding pattern, in which tandem repetitions may play a role in regulating oligomerization and macromolecular aggregate assembly. Because the in vitro-generated assemblies could represent only a fraction of the structures adopted in vivo, the similar aggregation propensity of the proteins into β-rich filaments supports the hypothesis that such interactions could promote in vivo structuring.

These findings complement previously conducted structural studies on the amyloid-like TasA and CalY proteins^32^ and expand our knowledge of the coordinated assembly of these proteins into filaments in the extracellular matrix of *B. cereus*. Interestingly, CapP adds another level of structural complexity compared to the orthologous pair TasA and CalY because it contains a C-domain region that is not involved in the rigid core but could still mediates CalY polymerization. Moreover, CapP (Figure S5B), similar to TasA and CalY,^32^ is disordered in vitro prior to aggregation, a notable difference from the TasA-TapA system of *B. subtilis*, which adopts a folded or partially folded conformation before assembly into fibrils.^30,32,61^ These distinctions suggest that this mechanism in *B. cereus* depends on an external trigger, which could be chemical or biophysical, such as the specific action of accessory proteins. This additional requirement has been previously proposed for the *in vivo* polymerization of CalY, and moreover, CalY has been described to enhance the polymerization of TasA without perturbing the global filament architecture.^22,32^ Based on these observations and our findings, it is reasonable to propose that the three proteins cooperate in assembling external pili and, consequently, organizing the ECM.

The diverse mechanisms that govern the assembly and attachment of non-flagellar proteinaceous filaments to bacterial cell surfaces include various structures, such as chaperone-usher pili, type V pili, type IV pili, curli, fap fibers, conjugative, type IV secretion pili, e-pili in diderm bacteria, and sortase-mediated pili as well as type IV pili in monoderm bacteria.^62,63^ In the monoderm *B. cereus*, BcpA-BcpB fimbriae are covalently bound to vegetative cells by sortase, while Ena appendages, exclusively found on spores, are assembled through subunit crosslinking via disulfide bonding.^64–66^ Nevertheless, our work describes a new type of fibrillar assembly exclusive to the *B. cereus* group, orchestrated by the chaperone function of CapP, with TasA and CalY serving as principal subunits. The biphasic effect, dependent on the concentration of CapP in CalY polymerization, has been reported for several extracellular chaperones.^67^ Under conditions where its levels are lower relative to the substrate, the chaperone acts to stabilize the conformation and promote filament formation. Conversely, at high concentrations, the chaperone forms soluble complexes with the substrate, completely inhibiting its assembly. Finally, it is noteworthy that disordered regions in chaperones are critical for interacting with target proteins.^68–70^ Although we have demonstrated the involvement of the C-domain of CapP in CalY polymerization, additional investigation is required to elucidate their precise structural and molecular contacts.

Building upon the novel insights unveiled in this study and the current knowledge on pili polymerization, we propose a revised model for the assembly of biofilm-specific pili in *B. cereus* (see Graphical abstract), which involves the paralogous proteins TasA and CalY, as well as the biofilm-specific protein CapP. Once the switch from a motile to sessile lifestyle has occurred, further growth and maturation of the biofilm ECM rely upon several factors, including the products of the operon *sipW-tasA* and the loci *capP* and *calY*, which are expressed under the tight control of the master regulator SinR. TasA, CapP and CalY are processed by the cognate signal peptidase SipW and secreted via the Sec pathway in an unfolded conformation. At 24 hours, the relative levels of CapP compared to CalY and TasA are lower, and associated with the cell surface, CapP interacts with unfolded CalY, catalizing the structural rearrangement necessary to initiate fibrillar polymer growth. Subsequently, CalY facilitates the incorporation of TasA monomers into growing pili. When the biofilm reaches a maturation stage at 48 hours, CapP levels rise, and it interacts with CalY within the ECM to form stable chaperone-substrate complexes that arrest the elongation of TasA-CalY heteropolymers. The ECM scaffold, composed of TasA-CalY filaments, exopolysaccharides, and eDNA, will provide *B. cereus* with the chemical and physical properties necessary for the correct assembly of the microbial community upon adhesion to biotic or abiotic surfaces.

Our findings contribute to knowledge on the pivotal role of the biofilm-specific protein CapP in orchestrating the formation and regulation of biofilms on *B. cereus* strains; in addition, the results provide insights into a new type of pilus assembly mechanism in monoderm bacteria. These remarkable advancements carry substantial implications for the clinical and food industries, in which innovative and effective strategies are urgently needed to address the critical challenge of *B. cereus* biofilm formation and its associated risks.

## ACKNOWLEDGMENTS

We thank Saray Morales Rojas for the technical support. We thank John Pearson from the Nanoimaging Unit of the Plataforma Bionand, Instituto de Investigación Biomédica de Málaga (IBIMA), for his technical support in confocal microscopy, data analysis and sample preparation. We thank Juan Félix López from the Nanoimaging Unit of the Plataforma Bionand, Instituto de Investigación Biomédica de Málaga (IBIMA), for his technical assistance in transmission electron microscopy imaging and analysis. We also thank Josefa Gómez Maldonado from the Ultrasequencing Unit of the SCBI-UMA for RNA sequencing and Luis Díaz Martínez from the SCBI-UMA for the transcriptomic analysis. The authors thank Carolina Lobo García and Casimiro Cárdenas García from the Proteomic Unit of the SCAI-UMA for protein sequencing and MS analysis. We also thank José Luis Zafra Paredes and Jose María Montenegro Martos from the Vibrational Spectroscopy Unit and the Electronic Spectroscopy Unit, respectively, of the SCAI-UMA for their technical support with the ATR-FITR spectroscopy and dynamic light scattering experiments. We are grateful to Theresa M. Koehler (University of Texas Health Science Center at Houston) for kindly providing the plasmid pUTE657 and to Stephen Leppla (National Institute of Health) for kindly providing us with the rat anti-sigmaA.

This work was supported by grants from an ERC Starting Grant (BacBio 637971), the Plan Nacional de I+D+i of the Ministerio de Ciencia e Innovación (PID2019-107724GB-I00 and PID2022-141664NB-I00), and Junta de Andalucía (P20_00479). Ana Álvarez-Mena is the recipient of an FPI contract (BES-2017-081275). This work benefited from the support of the French National Research Agency ANR (BH, grant No. ANR-23-CE11-0005-01), the Biophysical and Structural Chemistry Platform at IECB, CNRS UAR 3033, and INSERM US001.

## AUTHOR CONTRIBUTIONS

D.R. conceived the study. D.R., A.L. and A.A.M. designed the experiments and method analysis. A.A.M. performed the main experimental work and analyzed the data, with the following exceptions: A.A.M. and M.B.A.S. analyzed the SS-NMR data; M.L.A.G. performed some microbiological experiments; A.A.M., M.B. and B.K. performed biophysical analysis; A.G., E.M., and A.L. performed NMR analysis. JCA contributed to setting the initial hypothesis and performing iTRAQ experiments. A.A.M., M.B.A.S., B.H., A.L. and D.R. prepared the figures. A.A.M., B.H., A.A.L. and D.R. wrote the manuscript. A.V. contributed critically to writing the final version of the manuscript.

## SUPPLEMENTAL INFORMATION TITLES AND LEGENDS

**Figure S1.**
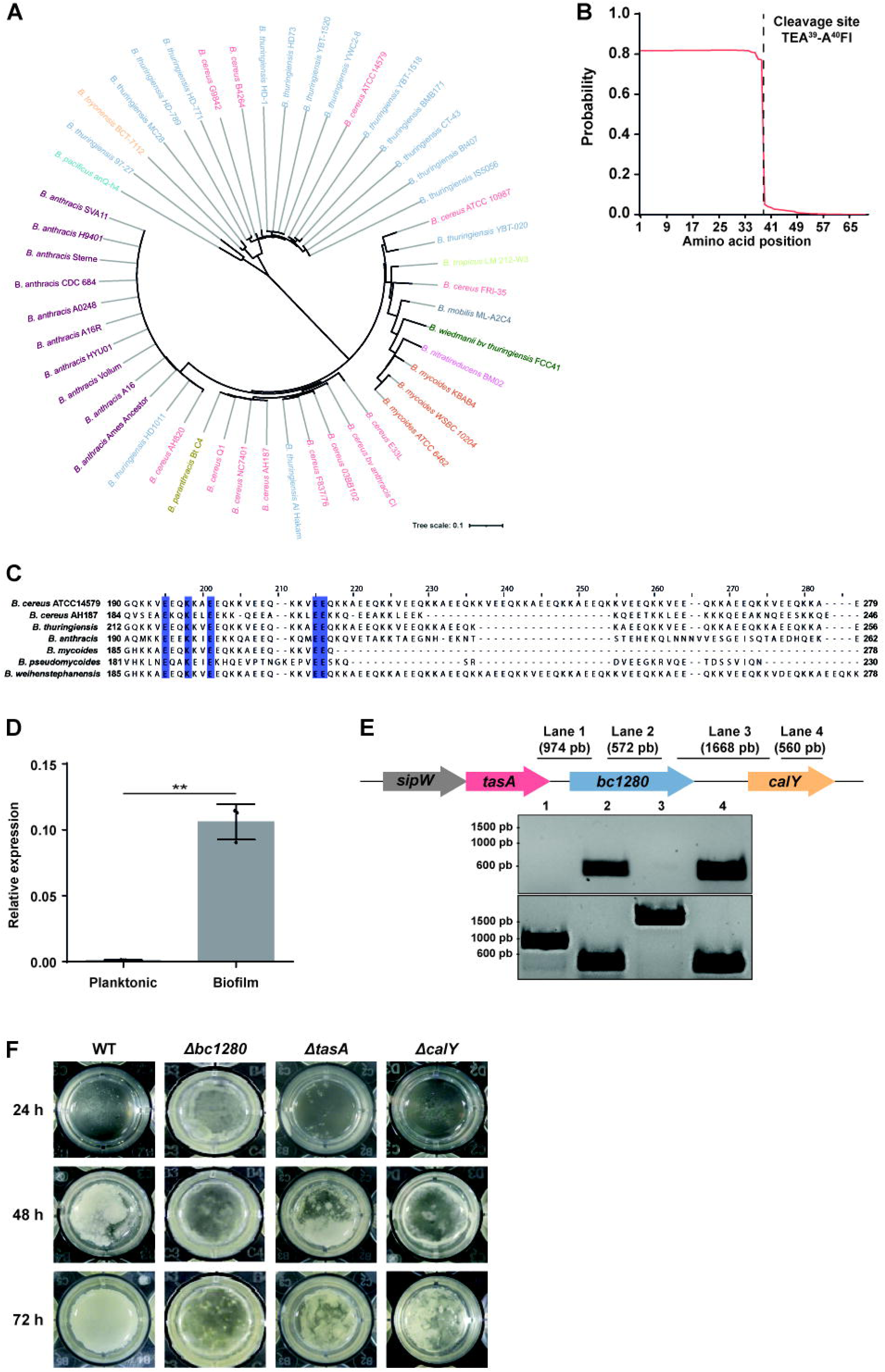
*bc1280* is exclusively found in the *B. cereus* group and is essential for biofilm formation. Related to Figure 1. **A)** Phylogenetic conservation of BC1280 among *B. cereus* group strains. Phylogenetic analysis was conducted on *B. cereus* group strains using the amino acid sequence of BC1280 (accession number in the KEGG database: BC1280) from *B. cereus* ATCC14579 as the reference. Evolutionary distances were calculated using the neighbor joining method. **B)** Signal peptide prediction by SignalP 5.0. A potential cleavage site was identified for the BC1280 protein between amino acid residues A^39^ and A^40^. **C)** Amino acid alignment of the C-domain among different strains of the *B. cereus* group, with color indicating conservation level. **D)** The relative expression of *bc1280* in planktonic and biofilm cells of the wild-type strain at 48 hours is presented. The average values from three biological replicates are displayed, with error bars indicating the standard deviation. Statistical analysis was performed using a two-tailed t test, revealing a significant difference between planktonic and biofilm cells (** p value=0.0055). **E)** Top panel: Bands corresponding to the products of amplification using cDNA from RNA extracted from the wild-type strain grown for 48 hours as a template are shown. Bottom panel: To confirm the functionality of the primers, each fragment was amplified via PCR using genomic DNA from the wild-type strain *B. cereus* ATCC14579 as a template. **F)** Biofilm phenotypes formed at the air-liquid interphase by the emetic strain *B. cereus* AH187 wild-type and the *tasA*, *calY* and *bc1280* mutant strains. Biofilm formation was evaluated as the formation of a pellicle at the air-liquid interphase after 3 days of incubation at 28°C.

**Figure S2.**
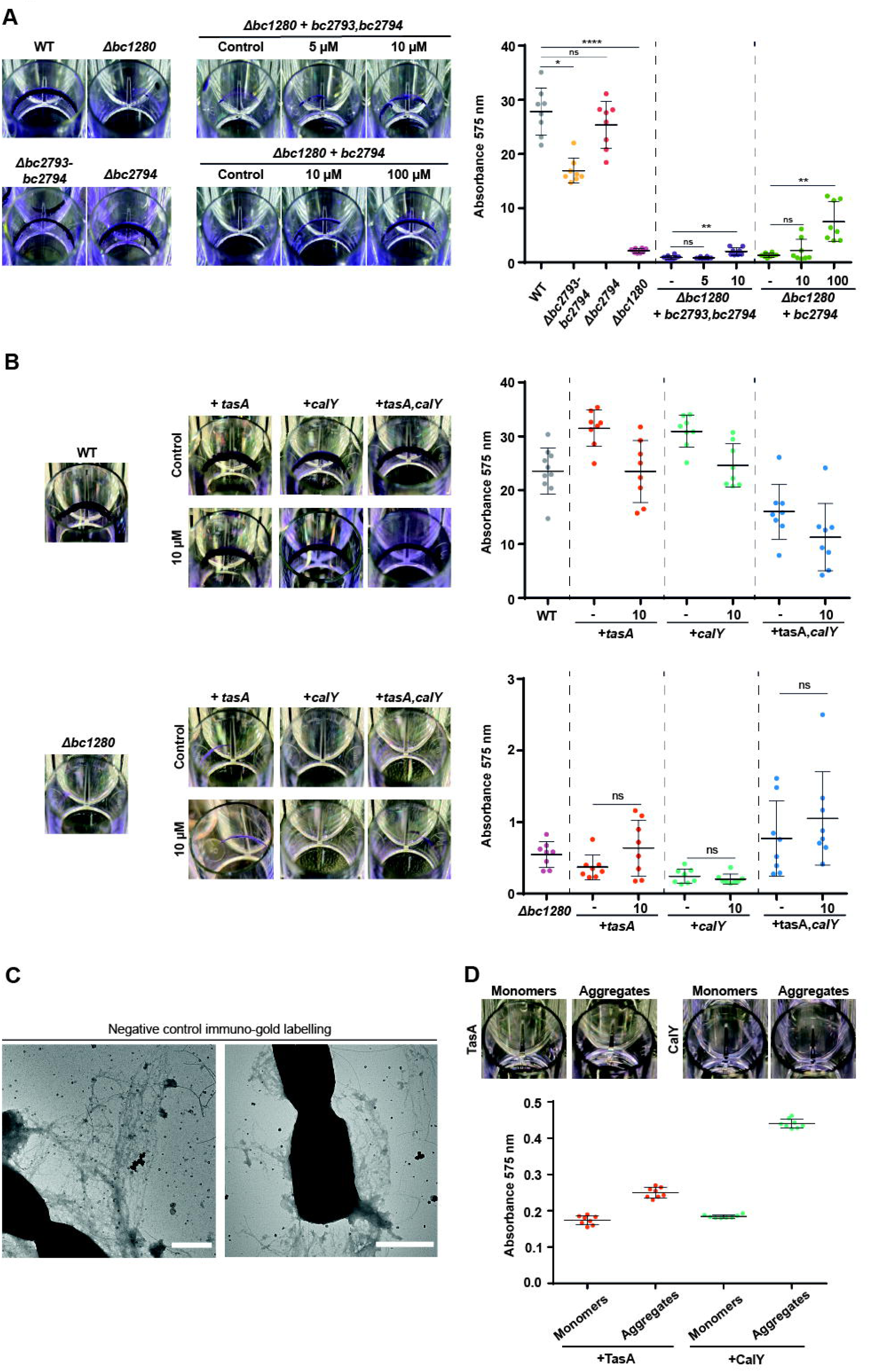
The genes *tasA*, *calY*, and an uncharacterized ECF sigma factor are deregulated in the *Δbc1280* mutant at 48 hours. Related to Figure 2**. A)** Biofilm rings formed after 72 hours by overexpressing *bc2793*-*bc2794* or *bc2794* in *Δbc1280* at various IPTG concentrations, as well as in *Δbc2793-bc2794* or *Δbc2794* strains, with wild-type and *Δbc1280* strains serving as controls. The biofilms were stained with crystal violet, and the biomass was measured by absorbance at 575 nm. Statistical significance was estimated using one-way ANOVA with Dunn’s multiple comparisons test (p < 0.05 (*), p < 0.01 (**), p < 0.001 (***), and p < 0.0001 (****). **B)** Biofilm phenotypes of the wild-type and *Δbc1280* mutant strains were assessed after overexpressing *tasA*, *calY*, or both genes using the plasmid pUTE657 with 10 μM IPTG for induction. As a negative control, each strain containing the plasmid was grown without IPTG. Biofilm formation was visualized by crystal violet staining. The biofilm mass was estimated by absorbance at 575 nm, and statistical significance was assessed by Dunn’s multiple comparisons test. No significant differences were detected between the control strain and upon induction with IPTG. **C)** Evaluation of the specificity of the nanogold particles in immunolabeling studies was conducted using the wild-type strain, with any primary antibody and the goat-antirabbit secondary antibody conjugated with 20 nm nanogolds (1:100). Scale bar: 1 μm **D)** Negative control showing TasA and CalY at 6 µM in monomer and fibrillar forms, grown in TY medium without bacterial inoculation. The wells were stained with crystal violet, and absorbance at 575 nm was measured to assess biofilm formation.

**Figure S3.**
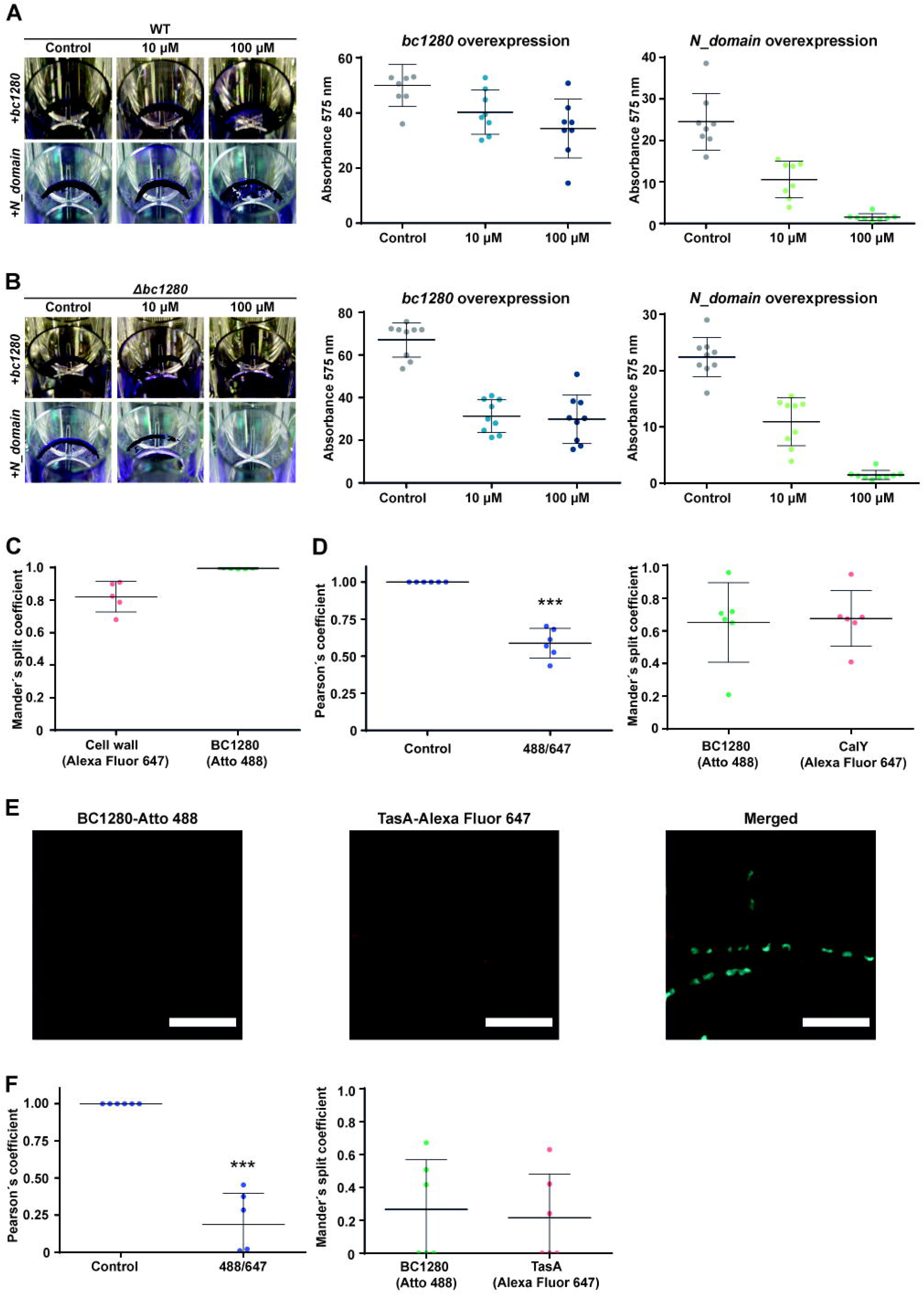
Biofilm formation is reverted upon overexpression of *bc1280* or *N-domain39-190* in *Δbc1280*, and BC1280 colocalizes with CalY but not with TasA. Related to Figure 3. **A)** and **B)** Overexpression of *bc1280* or the *N-domain* in the wild-type and *Δbc1280* strain at 10 and 100 µM IPTG. The resulting biofilms were stained with crystal violet and the absorbance at 575 nm was measured. **C)** Estimation of the proportion of fluorescence from co-localized pixels in each channel was performed to localize BC1280 in the cell wall using the Mander’s split coefficient and from five different fields of view. **D)** Pearson’s coefficient (*** p value = 0.0002 using a two-tailed t-test with Welch’s correction) and Mander’s split coefficient values obtained from BC1280-CalY CLSM studies. **E)** For colocalization studies of BC1280 and TasA using CLSM, a specific primary anti-TasA antibody (1:100) was utilized. Scale bar: 5 μm. **F)** Pearson’s coefficient (*** p value = 0.0002 using a two-tailed t-test with Welch’s correction) and Mander’s split coefficient values derived from BC1280-CalY CLSM studies.

**Figure S4.**
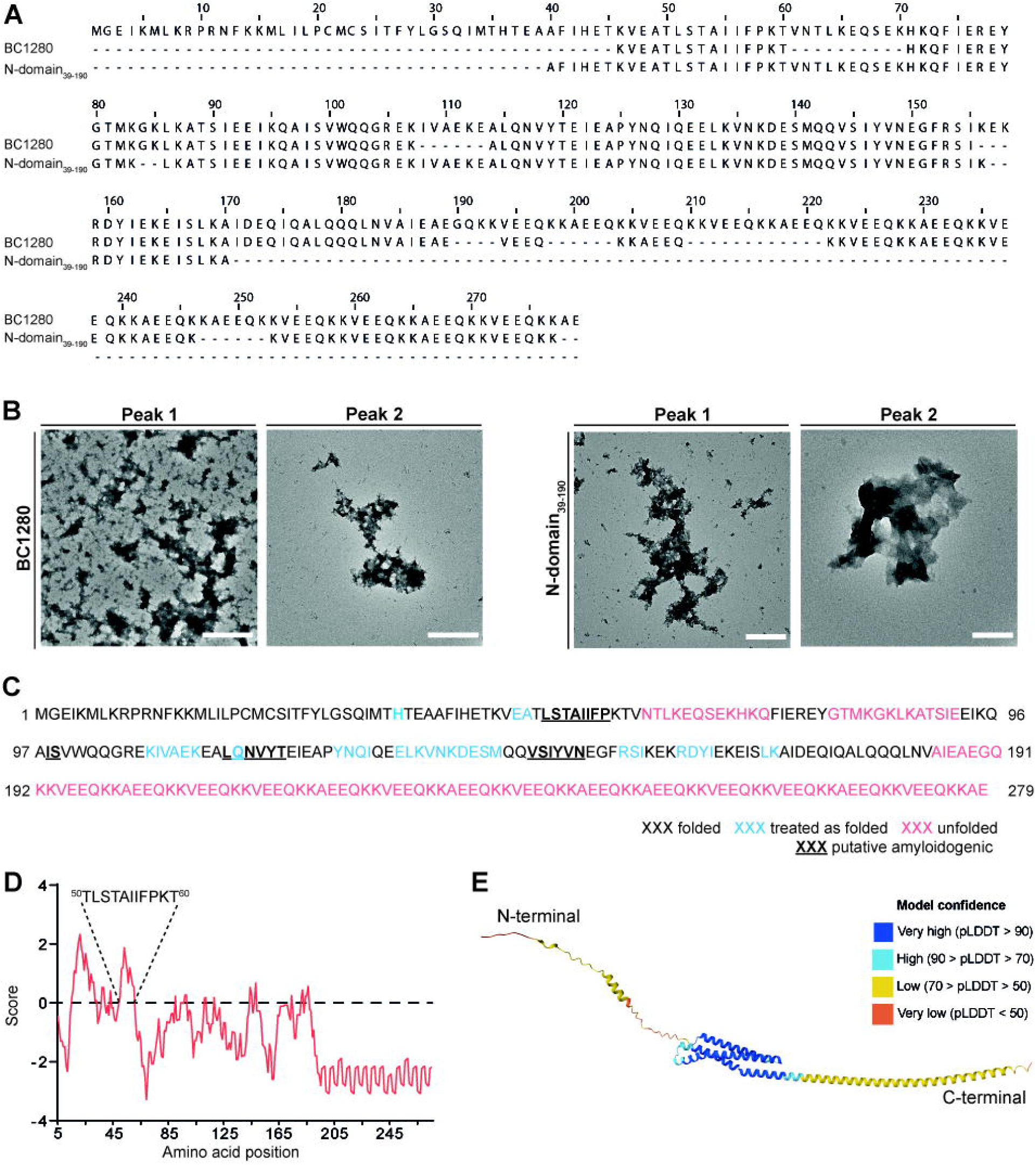
Preliminary studies indicate that BC1280 in vitro has a high tendency to oligomerize, and only the N-domain appears to be folded. Related to Figure 4. **A)** Mass spectrometry results obtained from the bands that corresponded to the elution fractions of BC1280 and N-domain39-190. **B)** Negative-staining micrographs of the first and second peaks detected by SEC for BC1280 and N-domain39-190. **C)** Prediction of the folding state using FoldUnfold software. The amino acid residues are classified as folded (black), folded (blue), or unfolded (red). The underlined amino acids are the regions identified as putative amyloidogenic by at least two different algorithms. **D)** Hydrophobicity prediction of residues in BC1280 using ProtScale software on the ExPASy server. **E)** The PDB model was constructed with the AlphaFold package for BC1280. The color pattern is displayed based on the confidence value.

**Figure S5.**
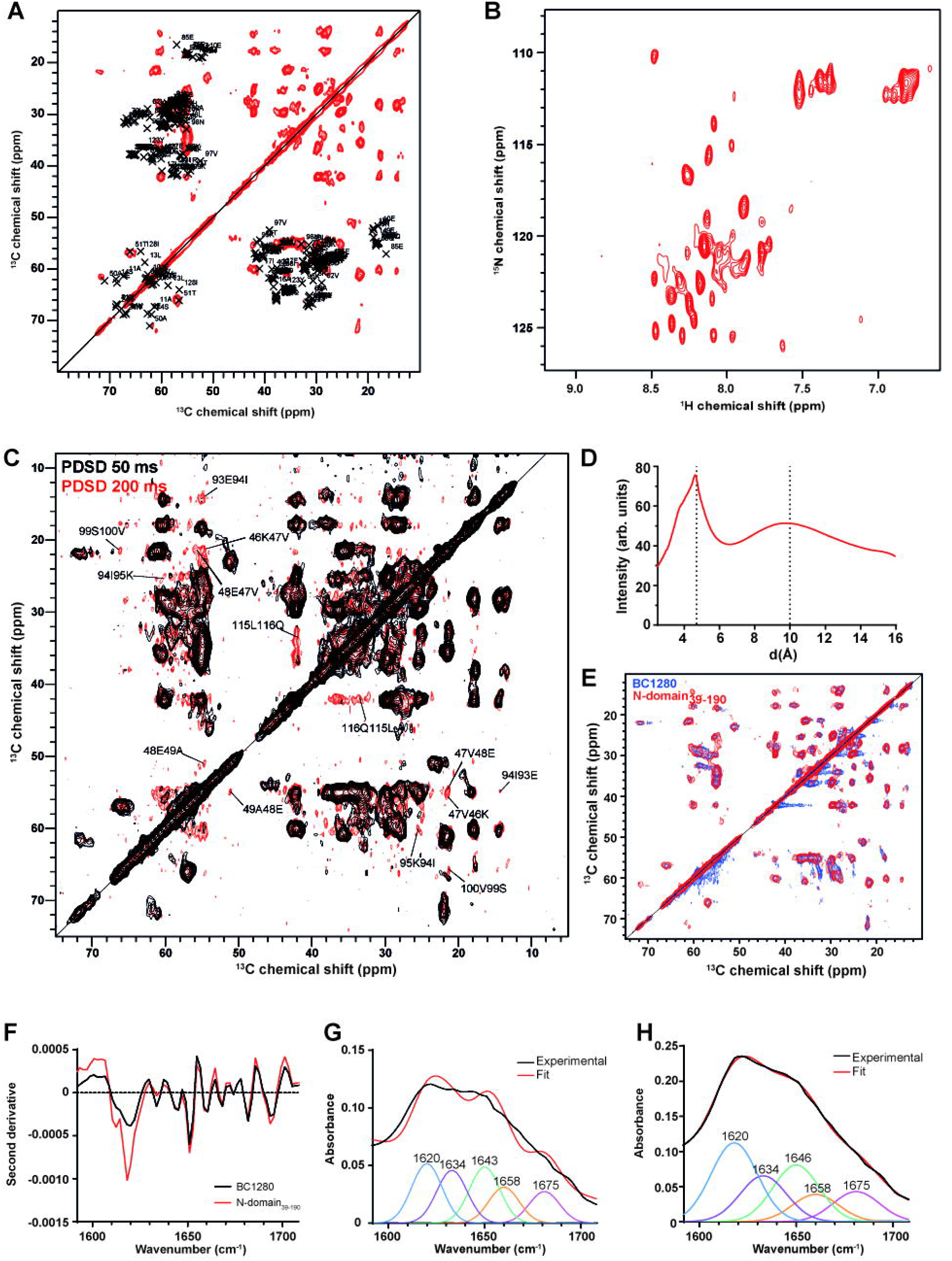
The structural rearrangement of the N-domain is conserved in the full-length protein. Related to Figure 5. **A)** Peaks predicted by SPARTA, based on the structure predicted by AlphaFold, are represented by the cross. The red signal indicates the experimental data obtained by SSNMR at a mixing time of 50 ms for the N-domain39-190. **B)** Solution NMR ^15^N-^1^H spectrum of N-domain39-190 protein demonstrates limited chemical shift dispersion. **C)** Cross-peak identification for inter-residue interaction. Overlay of the 2D ^13^C-^13^C PDSD correlation spectra obtained by SSNMR for N-domain39- 190 at 50 ms (black signals) and 200 ms (red signals) mixing times. The cross-peaks that correspond to chemical shifts between residues are marked. **D)** X-ray diffraction signals of BC1280 within 0–15 Å. **E)** Superimposition of the 2D ^13^C-^13^C PDSD correlation spectra obtained by SSNMR for BC1280 (blue) and N-domain39-190 (red) at a mixing time of 50 ms. **F)** Second derivative analysis of BC1280 (black) and N-domain39-190 in the amide I band using ATR-FTIR. **G)** and **H)** ATR-FTIR spectra of BC1280 and N-domain39-190, respectively, in the amide I band. The contribution of each peak was determined through secondary derivative analysis of the raw data, followed by a deconvolution approach.

**Figure S6.**
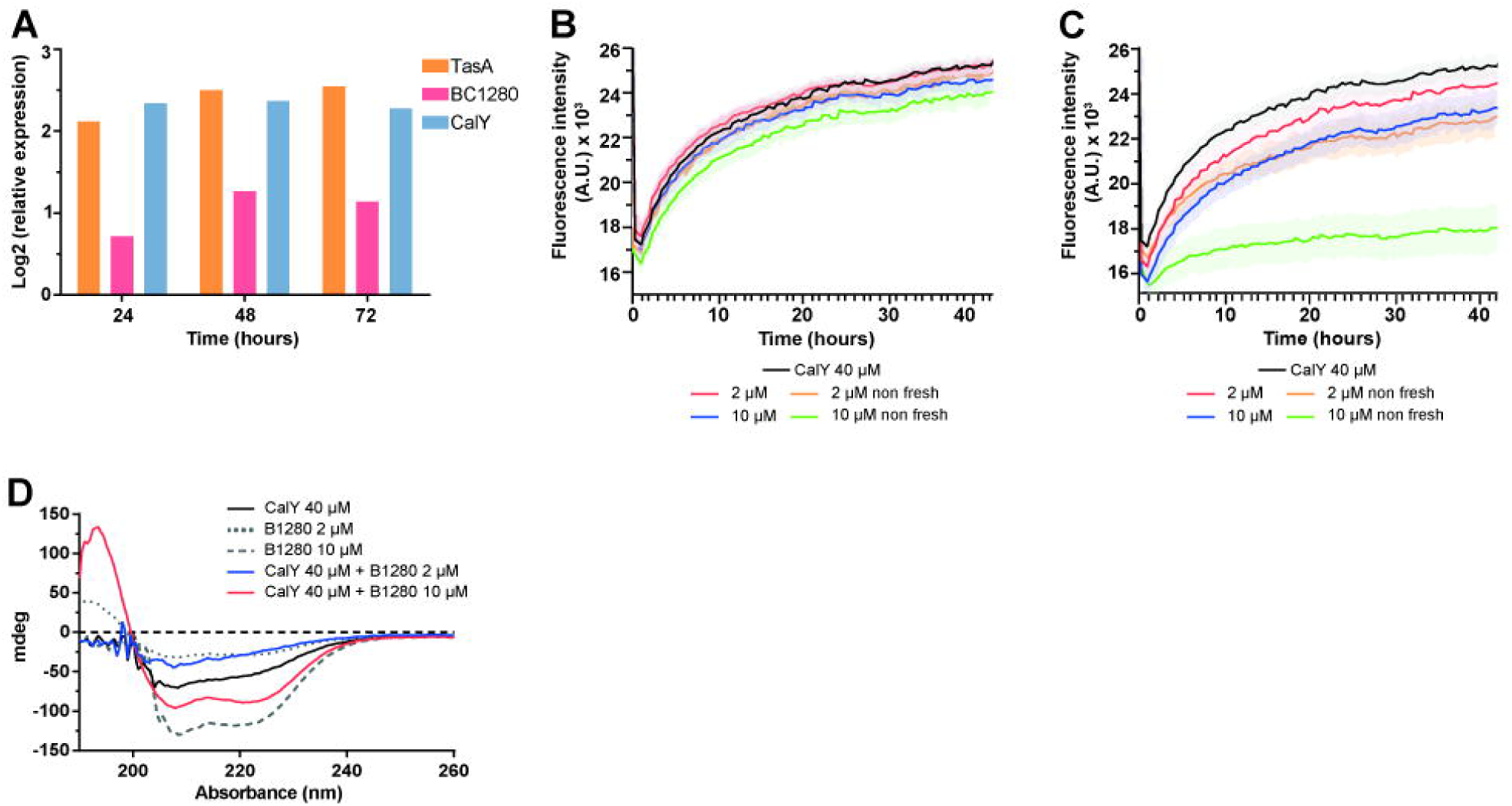
BC1280 levels have an impact on CalY polymerization. Related to Figure 6. **A)** Estimation of the relative levels of TasA, BC1280, and CalY in biofilm cells over 72 hours using iTRAQ. **B)** and **C)** Study of the effect of BC1280 and the N-domain39-190 without recent purification on CalY polymerization. ThT binding kinetics of CalY in the presence of freshly and non-freshly purified BC1280 (B) or N-domain39-190 (C) samples. Error bars indicate the standard error of the mean. **D)** Circular dichroism (CD) spectra obtained for each sample without subtracting the contribution of BC1280 at 2 and 10 μM when incubated with CalY for 16 hours under agitation.

## STAR METHODS

### RESOURCE AVAILABILITY

#### Lead contact

Further information and requests for resources and reagents should be directed to and will be fulfilled by the lead contact, Diego Romero.

#### Materials availability

Plasmids generated in this study are available at the BacBio Lab. Please contact Diego Romero for access. This study did not generate new unique reagents.

#### Data and code availability

RNaseq data have been deposited at GEO database and are publicly available as of the date of publication. Accession numbers are listed in the Key Resources Table. The mass spectrometry proteomics data have been deposited to the ProteomeXchange Consortium via the PRIDE partner repository with the dataset identifier PXD010211. Microscopy data reported in this paper will be shared by the lead contact upon request.

This paper does not report original code.

Any additional information required to reanalyze the data reported in this paper is available from the lead contact upon request.

### EXPERIMENTAL MODEL AND STUDENT PARTICIPANT DETAILS

#### Bacterial strains and culture conditions

The bacterial strains used in this study are listed in the Key Resources Table. For experiments conducted using *B. cereus* wild-type or mutant strains, cultures were routinely grown at 28°C from frozen stocks on Luria–Bertani (LB) plates containing 1% tryptone (Oxoid), 0.5% yeast extract (Oxoid), 0.5% NaCl and 1.5% agar before the respective experiments were performed. Biofilm assays were performed in TY medium supplemented with 1% tryptone (Oxoid), 0.5% yeast extract (Oxoid), 0.5% NaCl, 10 mM MgSO4 and 1 mM MnSO4.

For cloning and plasmid replication, the *E. coli* DH5α strain was used. For recombinant protein expression and purification, *E. coli* BL21(DE3) pLysS (Merck, Kenilworth, NJ, USA) was used. All the strains were cultured in Luria–Bertani (LB) liquid media. For the agar-solid plates, LB media was supplemented with 1.5% bacteriological agar (Oxoid). The final antibiotic concentrations were 100 μg/ml ampicillin, 50 μg/ml kanamycin, 100 μg/ml spectinomycin, and 5 μg/ml erythromycin.

For the purification of ^13^C/^15^N-labeled proteins, the cultures were grown in M9 minimal medium (the final concentration was 48 mM Na2HPO4, 22 mM KH2PO4, 8.6 mM NaCl, 1 mM MgSO4, 0.01 mM ZnCl2, 0.001 mM FeCl3, 0.1 mM CaCl2 and 10 ml 100X minimum essential medium vitamin solution) supplemented with 1 g/L ^15^NHCl as nitrogen and 2 g/L ^13^C6-D-glucose or 2-^13^C glycerol as the carbon source.

### METHOD DETAILS

#### Plasmid and strain construction

All the primers used to construct the strains used in this study are listed in Table S4.

*B. cereus* mutant strains were constructed via homologous recombination using the upstream and downstream regions of the gene of interest, which were subsequently cloned and inserted into the pMAD plasmid.^71^ The constructs were designed using NEB Builder HiFi DNA Assembly Master Mix (New England Biolabs, MA, USA) following the manufacturer’s instructions and utilizing specific primers. The pMAD vector was linearized using the restriction enzyme SmaI (FastDigest, Thermo Fisher Scientific), and the digested vector was incubated at 50°C for 1 hour along with the respective upstream and downstream fragments. The combined total amount of fragments and plasmid was 0.2 pmol, and a proportion of 1:2 (vector: fragments) was maintained. The resulting reaction mixture was subsequently transformed into *E. coli* DH5α, and positive colonies were selected using ampicillin. Subsequently, the plasmids were purified, subjected to PCR amplification and digestion for verification, and sequenced.

For the overexpression experiments in *B. cereus,* the gene of interest was cloned and inserted into the plasmid pUTE657^72^ under a promoter that was inducible by IPTG using the primers listed in Table S4. Following gene amplification and the inclusion of a specific RBS for *Bacillus*, the resulting PCR product was digested with SalI and SphI (FastDigest, Thermo Fisher Scientific) and subsequently cloned and inserted into the pUTE657 plasmid that was cut with the same restriction enzymes using T4 DNA ligase (Thermo Fisher Scientific). Subsequently, the resulting reaction was transformed into *E. coli* DH5α, and colonies were selected with 100 µg/ml ampicillin. After plasmid purification, all the vectors were subjected to PCR amplification, digestion with restriction enzymes, and sequencing.

The vectors were transformed into *B. cereus* by electroporation following previously described methods with some modifications. A single colony from a pure culture of *B. cereus* was inoculated in 5 ml of LB medium and incubated under shaking conditions at 30°C for 16 hours. This culture was subsequently used to inoculate a flask containing 100 ml of LB, which was incubated at 30°C with agitation until an OD600nm of 0.3 was reached. The cell culture mixture was centrifuged (3000 rpm, 5 minutes, 4°C), and the resulting pellet was resuspended in 12 ml of electroporation buffer (0.5 mM K2HPO4, 0.5 mM KH2PO4, 0.5 mM MgCl2, and 272 mM saccharose). The suspension was centrifuged again and then resuspended in 250 µl of electroporation buffer. The electrocompetent cells were mixed with 0.5–1 µg of plasmid and incubated on ice for 10 minutes. Next, the mixture was electroporated in a 0.2 cm cuvette using a voltage of 1.4 kV, a capacitance of 25 µF, and a resistance of 200 Ω. After electroporation, the suspension was recovered and incubated with 1.5 ml of LB for 5 hours at 30°C with agitation. The resulting culture was plated onto LB plates supplemented with the corresponding antibiotic to select the colonies that had been successfully transformed with the vector. For integration of the deletion, *B. cereus* transformants were incubated without antibiotics at 40°C, facilitating the integration of the mutation into the genome through homologous recombination. Finally, the culture was plated on LB agar plates, and the colonies were picked onto LB agar plates supplemented with erythromycin. Colonies that did not grow on LB supplemented with the antibiotic were selected, and the mutation was confirmed through PCR amplification.

For the heterologous expression of BC1280, the N-domain39-190 and the C-domain191-279 in *E. coli,* the open reading frames containing the *bc1280* gene, the region from amino acids 39 to 190. The PCR products and the pET24b vector were digested with NdeI and XhoI (FastDigest, Thermo Fisher Scientific) and subsequently cloned using T4 DNA ligase (Thermo Fisher Scientific) following the instructions supplied by the manufacturer. The reaction mixtures were subsequently transformed into *E. coli* DH5α, after which the colonies were successfully transformed with 50 µg/ml kanamycin.

#### Phylogenetic tree

The amino acid sequence of BC1280 was obtained from the KEGG database using *B. cereus* ATCC14579 as a reference. BLASTP analysis was performed, and the top 50 hits were aligned using Clustal Omega.^73^ The phylogenetic tree was then constructed using neighbor joining distances.

#### RT‒qPCR

Quantitative reverse transcription PCR was used to estimate the transcription levels of i) *bc1280* in planktonic and biofilm cells of the wild-type strain; ii) *tasA*, *calY*, *eps1* and *eps2* in *Δb1280* compared to those in the wild-type; and iii) *tasA*, *calY*, *eps1* and *eps2* in *Δb1280* overexpressing *bc2794* or *bc2793-2794* compared to those in the wild-type. RT‒ qPCR was performed using an iCycler-iQ system (Bio-Rad) and Power SYBR Green Master Mix (Thermo Fisher Scientific). Primer3 software (https://primer3.ut.ee/) was used for primer design, maintaining the default parameters.^74^ First, 1 µg of total RNA was reverse transcribed into cDNA using SuperScript III reverse transcriptase (Invitrogen) following instructions provided by the manufacturer. The reactions were performed in triplicate in 96-well plates with a total volume of 20 µl. The RT‒qPCR cycle started with an initial denaturation at 95°C for 3 minutes, followed by 40 amplification cycles (95°C for 20 seconds, 56°C for 20 seconds, and 72°C for 30 seconds), and the final step was 95°C for 30 seconds. The target genes, including *bc1280*, which encodes the protein BC1280; *bc1279*, which encodes TasA (also known as spore coat-associated protein N); *bc1281*, which encodes CalY (annotated as cell envelope-bound metalloprotease, camelysin); *bc5263*, which encodes UDP-glucose 4-epimerase; *bc5268*, which encodes secreted polysaccharide polymerase; *bc5277*, which encodes tyrosine-protein kinase; *bc5279*, which encodes tyrosine-protein kinase; and *bc1583*, which encodes O-acetyl transferase, were amplified using the primer pairs provided in Table S5. Primer efficiency and amplification product specificity were assessed as previously described.^75^ The estimation of relative expression levels was calculated using the ΔΔCt threshold (Ct) method.^76^ The housekeeping gene *rpoA* was used as a reference for data normalization. The relative expression value was calculated as the difference between the qPCR threshold cycle (Ct) of the gene of interest and the Ct obtained for *rpoA* (ΔCt=Ct*target gene* – Ct*rpoA*). The results obtained for the genes of interest were normalized to the values obtained for the wild-type strain. The RT‒qPCR analyses were conducted three times, each with three independent biological replicates.

#### Determination of the transcriptional unit by PCR

To determine whether *bc1280* constitutes an operon with *tasA* or *calY*, primers were designed between the genes and inside each gene to amplify four specific fragments that were 974 bp, 572 bp, 1668 bp, and 560 bp in size (primers listed in Table S6). cDNA was used as the template, and PCR was performed using genomic DNA purified with the commercial JetFlex^TM^ Genomic DNA Purification Kit (Thermo Fisher Scientific, Bremen, Germany) as a positive control. The fragments were amplified by PCR using Phusion^TM^ High-Fidelity DNA Polymerase (Thermo Fisher Scientific) with the following program: initial denaturation at 98°C for 30 seconds, followed by a 30-cycle amplification program (98°C for 10 seconds, 56°C for 30 seconds, and 72°C for 1 minute), and a final extension step at 72°C for 5 minutes. The samples were loaded onto an agarose gel and compared to the fragments amplified from genomic DNA.

#### Biofilm formation and extracellular complementation assays

To study the biofilm formation phenotype for each strain or condition, bacterial cultures were grown in 24-well plates as previously described.^25^ The strains were subsequently grown on LB agar plates at 28°C for 24 hours. The bacterial mass was resuspended in 1 ml of TY medium, and the OD600 was adjusted to 1. Subsequently, 10 µl of the suspension was inoculated into each well containing 1 ml of TY medium. The plates were then incubated at 28°C without agitation for 72 hours.

For the extracellular complementation assay involving the addition of TasA or CalY as monomers or in their fibrillated states, the proteins were previously purified *in vitro* from inclusion bodies through heterologous expression in *E. coli* following a protocol established by El-Mammeri et al., in 2019.^32^ Subsequently, the proteins were dialyzed against a 20 mM Tris and 50 mM NaCl solution at pH 7.4, after which the protein concentrations were measured. The assembly of filaments was promoted at 37°C under agitation at 200 rpm for one week. TasA or CalY protein, in both its monomeric and fibrillated forms, was added at a final concentration of 6 μM to 24-well plates inoculated with the *Δbc1280* strain. These plates were then incubated without agitation at 28°C for 72 hours, after which the resulting biomass was stained with crystal violet.

#### Crystal violet adhesion assay

Bacterial adhesion to the well plate surface was quantitatively measured using crystal violet staining, as previously described.^77^ The biofilms were grown for 72 hours, after which the medium was removed. Then, 1 ml of 1% crystal violet was added to each well of the 24-well plate, and the cells were incubated for 15 minutes. The plates were washed multiple times with water and left to dry for 45 minutes. Next, the biomass that was formed in each well was resuspended in 1 ml of 50% acetic acid, and the absorbance at 575 nm was measured for each sample using a plate reader (Tecan Infinite M1000 Pro; Tecan, Männedorf, Switzerland). The statistical significance was evaluated using one-way ANOVA with Dunnett’s multiple comparisons test. P values less than 0.05 were considered to indicate statistical significance.

#### Auto-aggregation assay

The effect of *bc1280* and *C-domain* deletion on the aggregation rate of *B. cereus* was estimated using a kinetic method following a previously described protocol.^24^ The strains were inoculated into a flask containing 20 ml of TY medium and incubated overnight at 28°C with agitation. The OD600 was adjusted to 3 for a final volume of 10 ml, after which the supernatant was centrifuged (5000 rpm, 10 minutes, 20°C) for dilution. The samples were then incubated in vertical tubes under static conditions at room temperature for 24 hours, after which the OD600 was measured every hour at the air‒liquid interface.

#### RNA extraction

For RNA extraction, a previously described protocol was followed with several modifications.^26^ Bacterial strains were cultivated in 24-well plates to retrieve the desired population—either planktonic or biofilm cells—at a specific incubation time. For the biofilm cells, the supernatant was removed from each well, followed by three wash steps, and the biomass was subsequently resuspended in 1 ml of PBS. To minimize variability between replicates, each sample was prepared by extracting 500 µl from 8 distinct wells. The bacterial suspensions were then centrifuged at 12000 × g for 5 minutes, and the resulting pellets were frozen at -80°C for 30 minutes or until needed. The pellets were resuspended in 900 µl of TRI-Reagent (Merck), and the cells were disrupted with 0.1 mm beads using a TissueLyser (QIAGEN) for 3 minutes, followed by incubation at 55°C for 3 minutes. Then, 200 μl of chloroform was added to each tube, which was vortexed for 10 seconds and incubated at room temperature for 3 minutes. Finally, the samples were centrifuged at 12000 × g for 10 minutes at 4°C. The resulting aqueous phase was transferred to a new tube containing 500 µl of ice-cold isopropyl alcohol. The tubes were inverted a few times, incubated for 10 minutes at room temperature, and then centrifuged at 12000 × g for 10 minutes at 4°C. The supernatants were subsequently removed, and the pellets were washed with 1 ml of ice-cold 75% ethanol, followed by centrifugation (12000 × g, 5 minutes, 4°C). Briefly, the RNA pellets were dried and then resuspended in 50 µl of DEPC-treated water. The residual DNA was eliminated through treatment with rDNAse, which is included in the Nucleo-Spin RNA Plant Kit (Macherey-Nagel), following the instructions provided by the manufacturer. The quality and integrity of the total RNA were assessed using an Agilent 2100 bioanalyzer (Agilent Technologies) and gel electrophoresis.

#### Whole-transcriptome analysis

A RNA-Seq analysis was performed on *Δbc1280* planktonic cells in comparison to the wild-type after 24 and 48 hours of incubation at 28°C. RNA extraction was performed following a previously described protocol, and each sample corresponding to a specific condition was collected in duplicate. RNA sequencing was performed by the Omics Unit of Supercomputing and Bioinnovation Center (SCBI, University of Malaga, Spain). rRNA removal was achieved using the RiboZero rRNA Depletion Kit (Illumina, CA, USA), and subsequently, 100-bp single-end read libraries were prepared using the TruSeq Stranded Total RNA Kit (Illumina). Next, the libraries were sequenced using the NextSeq550 instrument (Illumina). To eliminate regions of low quality, ambiguity, and low complexity, as well as potential contamination, the raw reads were preprocessed using SeqTrimNext with the specific configuration parameters used for NGS technology.^78^ Subsequently, the clean reads were aligned and annotated using the *B. cereus* ATCC14579 genome (NC_004722.1) as the reference. This alignment was conducted with Bowtie2, resulting in BAM files that were subsequently sorted and indexed using the SAMtools v1.484 program.^79,80^ To calculate the read count for each gene, the script Sam2counts (https://github.com/vsbuffalo/sam2counts) was used to determine the number of uniquely mapped reads. The analysis of differentially expressed genes (DEGs) between *Δbc1280* and the wild-type strain was conducted using the R script DEgenes Hunter. This script employs a combined p value, utilizing Fisheŕs method to consider the nominal p values obtained from edgeR and DEseq2.^81,82^ The combined p value was adjusted using the Benjamini–Hochberg (BH) test, which employs the false discovery rate approach. The adjusted p value was subsequently used to rank all the DEGs obtained. A p value < 0.05 and a log2FoldChange < -1 or > 1 were considered to indicate statistical significance.

#### Transmission electron microscopy and immunochemistry studies

To investigate the location of the proteins within the ECM using transmission electron microscopy, we followed a protocol that was previously established and described, with several modifications.^25,29,32^ For TasA and CalY immunodetection, the strains were cultured in multi-well plates at 28°C for 72 hours. Alternatively, for the localization of BC1280 or the N-domain within *B. cereus* cells, the strains corresponding to *Δbc1280* containing the plasmids pUTE657-*bc1280-6xHis* or pUTE657-*N-domain-6xHis* were grown in multiwell plates with TY medium supplemented with a 10 µM IPTG solution. The biofilm mass was subsequently harvested in 1 ml of PBS after 48 hours.

Then, 20 µl of each sample was applied to a copper grid and incubated for 2 hours. The grids were subsequently washed with PBS and incubated for 5 minutes to remove any excess bacterial cells. The cells that adhered to the grid were fixed using a 2% paraformaldehyde solution diluted in PBS and incubated for 10 minutes. The grids were washed again with PBS for 5 minutes and then blocked for 30 minutes using Pierce^TM^ Protein-Free T20 (TBS) Blocking buffer (Thermo Scientific). Next, the samples were incubated for 2 hours with the primary antibody at a concentration of 1:100, which was diluted in blocking buffer. Specific primary antibodies against TasA and CalY were used to detect TasA and CalY, respectively. For the immunolocation of BC1280-His and N-domain-His, we employed the primary antibody anti-6-His produced in rabbits (Merck). Subsequently, the grids were washed twice with TBS-T, each time with a 5-minute incubation. The samples were incubated for 1 hour with a goat anti-rabbit 20-nm gold-conjugated secondary antibody (Ted Pella, Redding, CA, USA) at a dilution of 1:100 in blocking buffer. Next, the grids were subjected to two additional washes with TBS-T, each lasting 5 minutes. The samples were fixed with 2.5% glutaraldehyde for 10 minutes. The samples were washed with Milli-Q water for 5 minutes and subsequently negatively stained with a 1% uranyl acetate solution. Finally, the grids were rinsed with a single drop of water and dried under dark conditions.

As a negative control to check the specificity of the 20-nm gold secondary antibody, we performed the same protocol, but instead of adding the primary antibody, we added blocking buffer at that step.

The samples were examined using a Thermo Fisher Scientific Tecnai G^2^ 20 TWIN transmission electron microscope at an accelerating voltage of 120 kV. The final images were taken using an Olympus Veleta side-mounted CCD with a resolution of 2k × 2k Mpx.

#### Cellular fractionation

For protein immunodetection by Western blotting, the samples were fractionated into three components: the extracellular medium, cell wall-associated proteins, and cellular contents (membrane and cytosol). Preliminary studies of BC1280 and the N-domain were conducted through cellular fractionation using the *Δbc1280* strain harboring the plasmid pUTE657-*bc1280*-6xHis or pUTE657-N*-domain*-6xHis, respectively. The strains were subsequently grown in TY media supplemented with agitation at 28°C. When the cultures reached an OD600nm of 0.4, 10 µM IPTG was added to each sample, and the cultures were incubated for 4 hours.

All the samples were subjected to the following protocol. The samples were centrifuged at 6000 × g for 5 minutes, after which the supernatant was passed through a 0.45 μm PES filter and retained as the extracellular medium fraction. The pellets were resuspended in 10 ml of PBS containing 100 µg/ml lysozyme and incubated for 2 hours at 37°C. Subsequently, the samples were centrifuged at 9000 × g for 20 min at 4°C, after which the resulting supernatant and the pellet were retained as the cell wall and cellular fractions, respectively. The cellular fraction was then resuspended in a small volume of PBS. Finally, all the samples were precipitated with trichloroacetic acid (TCA) to a final concentration of 10%. The mixture was then incubated on ice for 1 hour. Next, the precipitated proteins were centrifuged at 13000 × g for 20 minutes at 4°C, and the resulting pellets were washed twice with 1 ml of acetone. Finally, the samples were dried at 37°C for 5 minutes, resuspended in 1x Laemmli buffer (Bio-Rad), and loaded onto a 12% SDS‒PAGE gel for the immunodetection assay.

#### Polyacrylamide gels and western blotting

The samples were diluted in 1x Laemmli buffer (Bio-Rad) and then heated at 100 °C for 5 minutes. The proteins were separated by 12% SDS‒PAGE with the Spectra^TM^ molecular weight marker (Thermo Fisher Scientific) and subsequently transferred to a polyvinylidene fluoride (PVDF) membrane (Bio-Rad) using the Trans-Blot Turbo Transfer System (Bio-Rad) at 25 V for 30 minutes. Next, the membrane was blocked for 1 hour using 5% nonfat milk diluted in 50 mM Tris-HCl (150 mM NaCl, pH 7.5) containing 0.1% Tween-20 (TBS-T). The membrane was then incubated with the primary antibody in a solution of 3% nonfat milk in TBS-T. The membrane was washed three times with TBS-T, with 10 minutes of incubation between each wash. Next, the membrane was incubated with the secondary antibody against rabbit IgG conjugated to horseradish peroxidase (Bio-Rad) for 2 hours at a concentration of 1:3000 and diluted in TBS-T. Following the incubation, the membrane was washed twice with TBS-T and once with TBS. Finally, for immunodetection, the membranes were exposed to Pierce ECL Western blotting Substrate (Thermo Fisher Scientific).

For the immunodetection of BC1280-His or N-domain-His, we used an anti-6-His (Merck) primary antibody produced in rabbits at a dilution of 1:2500.

To study the molecular weight of the oligomers formed by BC1280 and N-domain39-190 under native conditions, the proteins were diluted in native sample buffer (Bio-Rad), loaded onto polyacrylamide gels (Any kD™ Mini-PROTEAN® TGX™ Precast Protein Gels, Bio-Rad), and analyzed using the NativeMark^TM^ (Invitrogen) marker as a reference. The native gels were run in buffer containing 25 mM Tris and 192 mM glycine at a constant voltage of 200 V. To visualize the proteins, the gels were stained with Coomassie brilliant blue.

#### Immunolocalization studies by fluorescence microscopy

For the immunofluorescence experiments, the samples were grown as previously described for transmission electron microscopy studies. Then, 150 µl of each sample was added to well slides coated with 0.1% poly-L-lysine (Sigma‒Aldrich) and incubated for 2 hours to facilitate bacterial adhesion to the substrate. The sample was then removed, and the bacteria that had attached to the slide surface were fixed with fixation buffer (3% paraformaldehyde and 0.1% glutaraldehyde diluted in PBS) for 10 minutes. Briefly, the wells were washed twice with PBS, and the samples were incubated for 1 hour with blocking buffer (3% m/v bovine serum albumin (BSA) and 0.2% v/v Triton X-100 in PBS). Then, the buffer was removed, and the wells were incubated for 3 hours with the primary antibody at a concentration of 1:100 diluted in blocking buffer. The primary antibodies used for visualization by TEM were also used for this experiment. The wells were then rinsed three times with washing buffer (0.2% m/v BSA and 0.05% v/v Triton X-100 in PBS), and each incubation lasted for 5 minutes. Next, the wells were incubated for 2 hours with the secondary antibody goat anti-rabbit IgG-Atto488 (manually labeled) at a dilution of 1:400 in blocking buffer. The samples were washed once with washing buffer and twice with PBS, for an incubation period of 5 minutes each. The immunostainings were fixed for 5 minutes using fixation buffer, and the wells were subsequently rinsed with PBS three times. To detect the cell wall, the slide was incubated with wheat germ agglutinin labeled with Alexa Fluor 647 (WGA, Thermo Fisher Scientific) for 20 minutes at a dilution of 1:100 in PBS. Finally, the bacterial DNA was stained for 20 minutes using Hoechst at a dilution of 1:1000. As a negative control, immunostaining was performed without incubation with the primary antibody.

The immunostaining was visualized using confocal laser scanning microscopy. For the fluorescence corresponding to Atto-488 visualization, we employed an excitation wavelength of 488 nm and detected the emission between 497 and 572 nm. The signal corresponding to Alexa Fluor 647 was visualized with an excitation wavelength of 561 nm, and emission was detected between 576 and 686 nm. The Hoescht signal was visualized independently using a specific dichroic filter.

For colocalization studies of BC1280-CalY/TasA, the same immunostaining protocol was followed with a few modifications. First, BC1280-6xHis was labeled using the anti-6-His antibody produced in rabbits (Merck), followed by goat anti-rabbit antibody conjugated to Atto-488 as the primary and secondary antibody, respectively. After three washes were performed with washing buffer, the samples were treated with quenching solution (0.1 M glycine in PBS) for 10 minutes. Next, TasA or CalY were detected using the following specific primary antibodies: anti-TasA and anti-CalY, which were produced in rabbit, followed by anti-rabbit conjugated to Alexa-Fluor 647 produced in goat as the secondary antibody. Following the final steps, the samples were fixed, stained with Hoechst, and finally visualized via CLSM, as described previously.

The colocalization images were analyzed using the software ImageJ and the colocalization threshold plugin. The region of interest (ROI) was defined for the region corresponding to Alexa Fluor 647, and both Pearsońs and Mandeŕs coefficients were calculated.^83^ As a negative control, the band corresponding to Alexa Fluor 647 overlapped with itself, resulting in 100% colocalization. For each sample, a minimum of 5 different images were analyzed, and statistical significance was determined for Pearsońs coefficient using an unpaired t test with Welch’s correlation, with a p value < 0.005.

#### Protein expression and purification

BC1280 and N-domain39-190 were purified under native conditions. For the heterologous expression of BC1280, the plasmid pET24b-*BC1280* was transformed into *E. coli* BL21 (DE3) (Merck, Kenilworth, NJ, USA), and colonies were selected with 50 µg/ml kanamycin. The plasmid pET24-*N-domain39-190* was subsequently transformed into *E. coli* Lemo21 (DE3)pLyss (New England Biolabs, USA) for N-domain39-190 expression, after which colonies were selected with 50 µg/ml kanamycin and 5 µg/ml chloramphenicol. The following steps were performed for both proteins. A single fresh colony was picked and cultured in 10 ml of LB supplemented with the corresponding antibiotics mentioned previously. The preculture was incubated under shaking conditions for 6 hours at 37°C. A total of 10% (v/v) of the precultures were inoculated into 1 L of LB supplemented with the corresponding antibiotics. The culture was incubated at 37°C with agitation until the optical density reached 0.6. For BC1280, the culture was induced with 400 µM IPTG and incubated for 16 hours at 28°C. Otherwise, the culture corresponding to the overexpression of N-domain39-190 was induced with 400 µM IPTG and 100 µM rhamnose and incubated for 3 hours at 28°C. Then, the cells were harvested (6000 × g, 30 minutes at 4°C, JLA 8.1 rotor, Beckman Coulter, Brea, CA, USA), and the pellets were frozen at - 80°C until use.

The pellets were resuspended in 18 ml of buffer A (20 mM Na2PO4, 500 mM NaCl and 20 mM imidazole, pH 8) with cOmplete™ EDTA-free Protease Inhibitor Cocktail (Roche) and CelLytic^TM^ B-Cell Lysis reagent (Merck). The lysates were incubated for 30 min with agitation at room temperature. The samples were sonicated using a Branson 450 digital sonifier on ice with 3 pulses for 45 seconds at 40% amplitude and then centrifuged at 15000 × g for 60 minutes at 4°C (F34-6-38 rotor; Eppendorf, Hamburg, Germany). The supernatant was loaded onto a HisTrap HP 5 ml column (GE Healthcare) and purified using an AKTA Start FPLC system (GE Healthcare). The column was preequilibrated with buffer A, and after sample loading, it was washed with the same buffer. Next, the proteins were eluted using a linear gradient of buffer B (20 mM Na2PO4, 500 mM NaCl and 500 mM imidazole, pH 8). Since the elution mixture was not completely pure, the sample was loaded onto a HiLoad 16/600 Superdex 75 pg column (Cytiva), and the pure protein was subsequently eluted in buffer containing 20 mM Tris and 50 mM NaCl, pH 7.5. The elution fractions were loaded onto a 12% Tris-Tricine SDS‒PAGE gel, and the samples containing the purified protein were concentrated using Amicon Ultra15 centrifugal filter units (Millipore) with a 3 kDa cutoff. Subsequently, the purified fractions were separated via 12% SDS‒PAGE to assess their purity, which was determined through Coomassie blue staining. Finally, the gel bands were analyzed using tandem mass spectrometry.

CalY and TasA were purified via heterologous expression in *E. coli* BL21(DE3) pLyss and from inclusion bodies under denaturing conditions following the protocol established by El Mammeri et al. ^32^

#### Tandem mass spectrometry analysis of protein bands

The SDS‒PAGE gel bands that corresponded to the protein purification and proteinase K digestion experiments were subjected to analysis via tandem mass spectrometry using a nanoion trap system [HPLC‒electrospray ionization–tandem mass spectrometry (MS)]. The bands were cut and destained using a mixture of 50% acetonitrile (ACN) and 25 mM ammonium bicarbonate. Then, the samples were dehydrated and dried using ACN. Disulfide bridges were reduced with 10 mM dithiothreitol (DTT) diluted in 50 mM ammonium bicarbonate, followed by incubation at 56°C for 30 minutes. Subsequently, the excess DTT was removed, and the cysteine residues were carbamidomethylated using 55 mM iodoacetamide diluted in 50 mM ammonium bicarbonate for 20 minutes at room temperature in the dark. The gel bands were then dehydrated again, and the proteins were digested using 10 ng/µl trypsin (Promega) at 30°C overnight. To extract the peptides from the gel, the samples were incubated with a solution of 0.1% ACN/formic acid (FA) for 30 minutes at room temperature. To eliminate ACN and ammonium bicarbonate, the samples were dried using a SpeedVacTM. Next, the samples were resuspended in a solution containing 0.1% FA, ultrasonicated for 3 minutes, and centrifuged at 13,000 × g for 5 minutes. Subsequently, the samples were purified and concentrated using C18 ZipTip® (Merck) following the instructions supplied by the manufacturer. Finally, the samples were injected into an Easy nLC 1200 UHPLC system, which was coupled to a Q Exactive HF-X Hybrid Quadrupole-Orbitrap mass spectrometer (Thermo Fisher Scientific). The software versions used for acquisition and analysis were Tune 2.9 and Xcalibur 4.1.31.9. The mobile phases used in the HPLC consisted of i) buffer A, which contained 0.1% FA dissolved in water, and ii) buffer B, which contained 0.1% FA dissolved in 80% ACN. Peptides were loaded onto a precolumn (Acclaim PepMap 100, 75 μm × 2 cm; C18, 3 μm; 100 A; Thermo Fisher Scientific) at a flow rate of 20 µl/min, while elution was performed using a 50 cm analytical column (PepMap RSLC C18, 2 µm; 100 A, 75 µm × 50 cm; Thermo Fisher Scientific). Elution was carried out through a gradient concentration over 60 minutes, transitioning from 5% to 20% buffer B. This was followed by a 5-minute gradient from 20% to 32% buffer B, resulting in a 10-minute elution with 95% buffer B. Then, the column was equilibrated with 5% buffer B using a constant flow rate of 300 µl/min. Before sample analysis, an external calibration was performed using LTQ Velos ESI Positive Ion Calibration Solution (Pierce, IL, USA), along with internal calibration using the polysiloxane ion signal at m/z 445.120024 obtained from ambient air. The MS1 scans were conducted within a m/z range of 375-1.600 at a resolution of 120.000. Employing a data-dependent acquisition strategy, the 15 most intense precursor ions with a charge ranging from +2 to +5 within a window of 1.2 m/z were selected from fragmentation to generate the corresponding MS2 spectra.

The fragmentation ions were generated through high-energy collision-induced dissociation with an initial mass set at 110 m/z and subsequently detected using a mass-analyzed Orbitrap at a resolution of 30.000. The dynamic exclusion for the selected ions was set to 30 seconds, and the maximum accumulation times for MS1 and MS2 were 50 ms and 70 ms, respectively. Finally, for protein sequence identification, the raw data were analyzed using Proteome Discoverer 2.4 (Thermo Fisher Scientific) with the Sequest HT search tool with mass tolerance parameters of 10 ppm and 0.02 Da for precursor and fragment ions, respectively.

#### Dynamic light scattering experiments

The samples corresponding to the elution peaks obtained by size exclusion chromatography for BC1280 and N-domain39-190 were filtered through a 0.46-µm syringe filter. The size measurements were performed using a Malvern Zetasizer Nano ZS (Malvern Panalytical, Malvern, United Kingdom) with a laser wavelength of 632.8 nm as the excitation source and a 1 cm pathway polystyrene cuvette as the sample holder. A total volume of 1 ml of the protein solution was transferred to the cuvette, and the dynamic light scattering signal was measured with a count rate ranging from 100 to 230 kcps, depending on the sample. Data collection and subsequent analysis were conducted using Zetasizer software v.6.34 (Malvern Panalytical).

#### Transmission electron microscopy

To study the morphology of BC1280, N-domain39-191 and C-domain191-279 purified *in vitro*, the samples were subjected to 2-fold serial dilutions and incubated on copper grids for 2 minutes. Then, the samples were stained with a 2% uranyl acetate solution for 1 minute and dried under dark conditions.

The samples were examined using a Thermo Fisher Scientific Tecnai G^2^ 20 TWIN transmission electron microscope at an accelerating voltage of 120 kV. The final images were captured using an Olympus Veleta side-mounted CCD with a resolution of 2k × 2k Mpx.

#### Thioflavin T (ThT) assays

The polymerization kinetics were studied using thioflavin T staining. The experiment was conducted in a 96-well plate, and 40 µM ThT was added to each well at a specific concentration, after which the mixture was diluted in buffer containing 20 mM Tris and 50 mM NaCl at pH 7.4. The polymerization kinetics of CalY were assayed at 40 µM using low concentrations of BC1280 and N-domain39-190 (2 and 10 µM).

The plates were incubated at 37°C in a plate reader and shaken at 100 rpm before each measurement (Tecan Infinite M1000 Pro; Tecan, Männedorf, Switzerland). The fluorescence intensity was monitored every 30 minutes through the bottom of the plate using an excitation wavelength of 440 nm and an emission wavelength of 480 nm. All measurements were recorded in triplicate.

#### Sequence analysis by different bioinformatics tools

The primary sequence of BC1280 was obtained from UniProt (https://www.uniprot.org/) with the accession number Q81GC7. SignalP5.0 was used to predict the cleavage site for the signal peptidase.^84^ The hydrophobicity was analyzed using the ProtScale tool provided by the ExPASy server, and the prediction of disordered regions was conducted using FoldUnfold with default parameters.^45,47^ Putative amyloidogenic regions were predicted employing the following bioinformatics tools: AmylPred2, FoldAmyloid, MetAmyl, PASTA2.0 and TANGO.^40–44^

#### AlphaFold prediction

The secondary structure model of BC1280 was predicted by AlphaFold using the sequence available in UniProt with the accession number Q81GC7 and default parameters.^48,49^ The pdb model obtained was subsequently used for predicting chemical shifts via SPARTA.^53^ Finally, the predicted correlations overlapped with the experimental data obtained by SSNMR.

#### *In vitro* assembly of oligomers

BC1280 and N-domain39-190 were diluted to a final concentration of 50 µM in buffer containing 20 mM Tris and 50 mM NaCl at pH 7.5. The oligomers self-assembled under agitation for 2 weeks at 37°C and a speed of 200 rpm. The aggregates were then ultracentrifuged (40k rpm speed, 5 hours, 18°C, TLA 120.1 rotor; Beckman Coulter), and the resulting pellets were analyzed using X-ray diffraction, ATR-FTIR, and SSNMR.

#### X-ray diffraction measurements

Filament diffraction patterns were obtained at 4°C using a Rigaku FR-X rotating anode X-ray generator (Rigaku, Tokyo, Japan) equipped with an EIGER 1 M hybrid pixel detector (Dectris, Baden, Switzerland) at the copper wavelength. The concentrated hydrated samples were mounted in MicroLoops from Mitegen (Ithaca, NY, USA) on a goniometer head under cold nitrogen flow. Each diffraction pattern represents a 360° rotation along the φ axis, with an exposure time of 720 s. No correction, such as smooth filtering or baseline correction, was applied to the data. WinPLOTR (https://cdifx.univ-rennes1.fr/winplotr/winplotr.htm) was used for converting q values to reticular distance (d) values before plotting.

#### Proteinase K (PK) digestion

A 22.5 µg solution of aggregated N-domain39-190 was treated for 45 minutes at 37°C with 3 µg/ml proteinase K in 20 mM Tris and 50 mM NaCl at pH 7.5. The proteinase activity was stopped using 5 mM PMSF. The samples were mixed with an equal volume of Laemmli buffer (Bio-Rad) and heated at 100°C for 5 minutes. Subsequently, the samples were separated using 12% SDS‒PAGE, and the bands were visualized by Coomassie blue staining. The bands of interest were excised and subjected to analysis by tandem mass spectrometry.

#### Attenuated total reflection FTIR spectroscopy

The samples were treated under the same conditions as those used for the X-ray studies. Infrared spectra for BC1280 and N-domain39-190 were recorded with an FT-IR spectrophotometer (model Vertex70; Bruker) in attenuated total reflectance (ATR) mode utilizing a Golden Gate Single Reflection Diamond ATR System accessory (Specac). No sample preparation was needed, and spectra were recorded with an air background and 64 scans for both the background and sample measurements. The spectral resolution was set at 4 cm-1”. The raw data within the spectral range of the amide I region were used for estimating the total amount of secondary structure. To increase the resolution of the minimal absorbance peaks, a second derivative was calculated. Next, a deconvolution approach was employed using PeakFit software to determine the contribution percentage for each type of secondary structure: parallel β-sheet (1634 cm^-1^), antiparallel β-sheet (1620 cm^-1^ and 1690 cm^-1^), α-helix (1658 cm^-1^), turns (1675 cm^-1^) and random coil (1646 cm^-1^), in accordance with the well-established assignments.^58^

#### Proteomic analysis by the isobaric tag for relative and absolute quantification (iTRAQ) method

To estimate the relative levels of TasA, BC1280 and CalY during biofilm formation in the wild-type strain *B. cereus* ATCC14579, we conducted sample preprocessing and subsequent analysis following the protocols described by Caro-Astorga et al.,in 2020.^26^

#### NMR spectroscopy

Solid-state NMR experiments were performed on a 600 MHz spectrometer (Bruker Biospin, Germany) equipped with a triple resonance 4mm probe. Experiments were performed at 11 kHz MAS frequency at a sample temperature of 283K. Chemical shifts were referenced with DSS. For 2D ^13^C-^13^C PDSD spectra (mixing time of 50 and 200 ms), an initial cross-polarization step of 0.5 ms was used and high-power SPINAL-64 decoupling was applied during acquisition times. 2D ^13^C-^13^C PDSD (50 ms) and ^1^H-^13^C INEPT spectra were recorded for 1-3 days, and 6 days for the ^13^C-^13^C PDSD spectrum (200 ms).

Solution NMR ^1^H,^15^N SOFAST-HMQC experiment was carried out at 298 K on a protein sample containing 200 μM BC1280, 20mM Tris, 50mM NaCl, pH7.5, on a 700 MHz spectrometer (Bruker Biospin, Germany) equipped with a 5mm TXI ^1^H/^13^C/^15^N/^2^H probe. The NMR data were processed using the TOPSPIN 4.0.6 software and analyzed by CCPNMR.^85^

#### Circular dichroism

CalY (40 μM) alone or in combination with BC1280 (at 2 or 10 μM) was incubated in 10 mM sodium phosphate buffer at pH 8 for 16 hours at 37°C under agitation. Circular dichroism spectra were recorded using a JASCO J-815 spectrometer, using a 0.1 cm path-length cuvette in the range of 180-260 nm, with a 0.5 nm step and a 1 s collection time per step. The scan rate was 50 nm/min. Final spectra were obtained as the average of six scans after a blank correction. The spectrometer was continuously purged with dry N2 gas.CD spectra were buffer subtracted, and the results were expressed as mean residue ellipticity.

